# Reduced EEG Complexity and Its Association with Social Communication in Adults with Autism Spectrum Disorder: A Multiscale Entropy Study

**DOI:** 10.64898/2026.09.01.747412

**Authors:** Taiga Naoe, Yumi Shikauchi, Takashi Itahashi, Kosuke Tanaka, Tsukasa Okimura, Ryu-ichiro Hashimoto, Haruhisa Ohta, Yosuke Sato, Motoaki Nakamura

**Affiliations:** Medical Institute of Developmental Disabilities Research, Showa Medical University, 6-11-11 Kita-Karasuyama, Setagaya-ku, Tokyo, 157-8577, Japan; Department of Linguistics, Graduate School of Arts and Letters, Tohoku University, 27-1 Kawauchi, Aoba-ku, Sendai, Miyagi 980-8576, Japan; Brain Function Analysis & Digital Medicine Research Institute, Showa Medical University, 1-5-8 Hatanodai, Shinagawa-ku, Tokyo, 142-8555, Japan; Department of Language Sciences, Graduate School of Humanities, Tokyo Metropolitan University, 1-1 Minami-Osawa, Hachioji-shi, Tokyo, 192-0397, Japan

**Keywords:** Autism spectrum disorder, Electroencephalography, Multiscale entropy, Neural complexity, Naturalistic stimuli, Resting state, Social communication, ADOS-2

## Abstract

**Background:** Brain functions emerge from temporally organized neural dynamics, and an appropriate level of neural complexity may support flexible information processing. Electroencephalographic (EEG) studies using multiscale entropy (MSE), which quantifies signal complexity across multiple temporal scales, have reported reduced MSE at longer scales in individuals with autism spectrum disorder (ASD). However, most evidence comes from studies of infant and child samples, leaving adult data scarce, and the associations between reduced MSE, clinical symptoms, and social information processing are insufficiently understood.

**Methods:** We recorded and analyzed EEG data from adults with ASD (n = 47) and typically developing (TD) controls (n = 40) during eyes-closed rest and two movie viewing conditions: *Inscapes*, comprising dynamically changing abstract visual patterns, and *Partly Cloudy*, an emotionally engaging animated social narrative. Group and condition effects were assessed using cluster-based permutation tests, and associations between MSE and Social Affect (SA) scores from the Autism Diagnostic Observation Schedule, Second Edition (ADOS-2), as well as group differences in event-related MSE changes, were examined using linear mixed-effects models.

**Results:** Across conditions, adults with ASD showed lower MSE at longer scales (τ = 16–30), corresponding to effective sampling rates of 12.50–6.67 Hz in the coarse-grained time series, than TD adults. Longer-scale MSE within the ASD–TD difference cluster was negatively associated with ADOS-2 SA scores across diagnostic groups. The difference in event-related MSE change between Social and Non-Social conditions, comparing empathic-pain and mentalizing events in *Partly Cloudy* with temporally matched non-social windows in *Inscapes*, was smaller in ASD participants than in TD participants. No significant difference between Social and Non-Social windows was observed among ASD participants, whereas TD participants showed significantly greater MSE change during Social than Non-Social windows.

**Limitations:** This study used a modest sample without an independent replication cohort, and the short event windows limited the precision of the MSE estimation.

**Conclusions:** Reduced longer-scale MSE extends to adults with ASD and is associated with higher ADOS-2 SA scores across diagnostic groups. Event-related findings further suggest attenuated differentiation of longer-scale MSE between socially relevant and matched non-social windows in ASD.

## Background

Autism spectrum disorder (ASD) is a neurodevelopmental condition characterized by persistent difficulties in social communication and interaction, along with restricted and repetitive patterns of behavior, interests, and activities [1, 2]. These clinical features are thought to involve atypical information processing, particularly in situations requiring flexible integration of social, sensory, and contextual information [3–5]. Conventional neuroimaging and electrophysiological approaches have provided important insights into atypical brain structure, activation, connectivity, and oscillatory activity in ASD [6–9]. At the same time, these approaches may not fully capture the temporal dynamics and complexity of neural activity that support flexible information processing across different timescales [10–13].

Electroencephalography (EEG) is well suited for characterizing neural dynamics because it captures neural activity with high temporal resolution. Prior EEG studies in ASD have primarily characterized oscillatory activity and interregional coordination, including functional connectivity and large-scale network organization [9, 14–18]. However, the direction and spatial distribution of these findings have varied across studies [9, 18–20], and robust EEG-based physiological markers of ASD have yet to be established. This heterogeneity partly reflects differences in participant developmental stages as well as variation in the recording context, such as eyes-open or eyes-closed rest, cognitive task engagement, and video viewing [19, 20]. In addition, many of these commonly used measures provide time-averaged summaries of neural activity and may therefore obscure aspects of the temporal organization of neural activity.

Several time-resolved EEG methods, including time–frequency analysis [21], dynamic functional connectivity [22], dynamic network organization [23], and transient phase-synchronization analysis [24], have been used to characterize temporal fluctuations in local neural activity and interregional coordination. Although these approaches provide valuable insights, they do not directly quantify the regularity and complexity of temporal patterns within local neural signals across multiple timescales. Neural complexity refers to the variability and richness of the temporal patterns of brain activity, reflecting the brain’s capacity to generate diverse yet organized states. An appropriate level of neural complexity may allow the brain to transition flexibly between states, thereby supporting adaptive information processing [25–27]. Such flexibility is particularly relevant to the processing of ambiguous information, which often requires the integration of multiple contextual cues [28, 29]. Social interactions provide a salient example, as others’ intentions and emotions must often be inferred from ambiguous and context-dependent signals.

Multiscale entropy (MSE) is a measure of nonlinear physiological signal complexity that quantifies temporal irregularity by estimating the sample entropy (SampEn) across multiple time scales [30, 31]. In contrast to various nonlinear measures that characterize signal complexity at a single scale or resolution [32], MSE provides an index of scale-dependent neural complexity. Previous EEG studies using MSE have identified altered neural complexity in several neuropsychiatric conditions [33–36]. In ASD, reduced MSE at longer temporal scales has been reported relative to healthy controls [5, 37–41], which is consistent with the proposal that ASD is characterized by reduced flexibility in information processing and integration [3–5]. Notably, this pattern contrasts with reports of atypically increased longer-scale MSE in schizophrenia and Alzheimer’s disease [33–35], raising the possibility that MSE is sensitive to differences in cortical dynamics and information integration across neuropsychiatric and neurodegenerative conditions [33, 34, 36]. Collectively, these findings suggest that the MSE may provide a physiological index of atypical neural dynamics in patients with ASD.

However, the existing EEG-MSE literature on ASD shares several limitations with the broader EEG literature [5, 37–41]. Most studies have focused on infants or children, with comparatively limited evidence in adults. Recording contexts have also varied substantially across studies, including eyes-open and eyes-closed rest and engagement in cognitive tasks. However, few studies have examined multiple experimental conditions within the same individuals, leaving unclear whether reduced MSE in ASD is consistently expressed across brain states or whether the magnitude and extent of group differences vary by condition. Moreover, the extent to which MSE is related to the core clinical features of ASD, particularly difficulties in social communication and the underlying processing of social information, has rarely been directly examined.

In the present study, we applied MSE to EEG data acquired from adults with ASD and typically developing (TD) adults under three within-participant experimental conditions: eyes-closed resting state and two naturalistic movie viewing conditions. The participants viewed two short animated films: *Inscapes* [42], which comprises dynamically changing abstract visual patterns, and *Partly Cloudy* [43], an emotionally engaging social narrative [44, 45]. Importantly, data-driven analyses have identified temporally extended segments of *Partly Cloudy* that reliably engage brain networks implicated in theory-of-mind (ToM) and empathic-pain processing [44, 45]. This feature made *Partly Cloudy* particularly well-suited to our aim of examining whether EEG dynamics expressed over multi-second time windows are related to clinically relevant individual variations in social communication characteristics. This naturalistic design enabled us to characterize MSE across conditions differing in cognitive and social demands while providing richer temporal and contextual information that more closely approximates aspects of real-world cognition than tightly controlled experimental tasks [44, 46, 47]. We further examined whether MSE was associated with individual variations in social communication difficulties, focusing specifically on the Social Affect (SA) subscore of the Autism Diagnostic Observation Schedule, Second Edition (ADOS-2) [48–50].

## Methods

### Ethics statements

All the participants provided written informed consent to participate in this study. The recruitment and experimental protocols were approved by the Institutional Review Boards of Showa Medical University (approval no. 2023-181-A) and Showa Medical University Karasuyama Hospital (approval no. B-2021-007), and the study was conducted in accordance with the Declaration of Helsinki.

### Participants

A total of 57 adults with neurodevelopmental disorders and 52 TD adults participated in an EEG experiment conducted between April 2024 and May 2026. All participants with neurodevelopmental disorders were outpatients at the Showa Medical University Karasuyama Hospital. Of these, 52 had received a formal diagnosis of ASD according to the Diagnostic and Statistical Manual of Mental Disorders, Fifth Edition (DSM-5) [1], or pervasive developmental disorder according to the DSM, Fourth Edition, Text Revision (DSM-IV-TR) [51].

TD individuals who voluntarily participated and had no history of neurodevelopmental or psychiatric disorders or major physical disabilities were enrolled as controls.

The participants completed ADOS-2 Module 4 [48, 49]. Intellectual ability was assessed using either the Wechsler Adult Intelligence Scale–Third Edition (WAIS-III) or the WAIS-Fourth Edition (WAIS-IV). The Intelligence Quotient scores were additionally estimated using the Japanese version of the National Adult Reading Test [52].

Of the 57 participants with neurodevelopmental disorders and the 52 TD participants initially enrolled, the following were excluded from the analysis: five participants with neurodevelopmental disorders who did not have an ASD diagnosis; three participants with ASD and one TD participant whose ADOS-2 assessments were unavailable; and two participants with ASD and seven TD participants with missing data for at least one of the three EEG conditions (resting or the two-movie viewing). The missing data resulted from experimental interruptions, incorrect montage settings during EEG recording, errors in recording the onset and offset triggers, or failures in data saving. In addition, three TD participants were excluded because their ADOS-2 scores exceeded the diagnostic threshold, and one TD participant was excluded because prior familiarity with psychological and clinical assessment procedures could not be ruled out, raising concerns about the validity of the WAIS and ADOS-2 scores.

The final analytical sample comprised 47 ASD and 40 TD participants (Table 1). Among these participants, 37 ASD and 39 TD participants completed the WAIS-IV, and 6 ASD participants completed the WAIS-III. The remaining four ASD participants and one TD participant did not complete the WAIS assessment; however, their intelligence quotient estimates based on the Japanese version of the National Adult Reading Test were within the normal range (115.76, 105.65, 107.67, 113.74, and 103.63). Among the 47 participants with ASD, 14 were taking at least one psychotropic or neurological medication, including antidepressants (n = 4), anxiolytics (n = 3), hypnotics/sleep medications (n = 4), antipsychotics (n = 6), and antiepileptic drugs (n = 5).

**Table 1.**
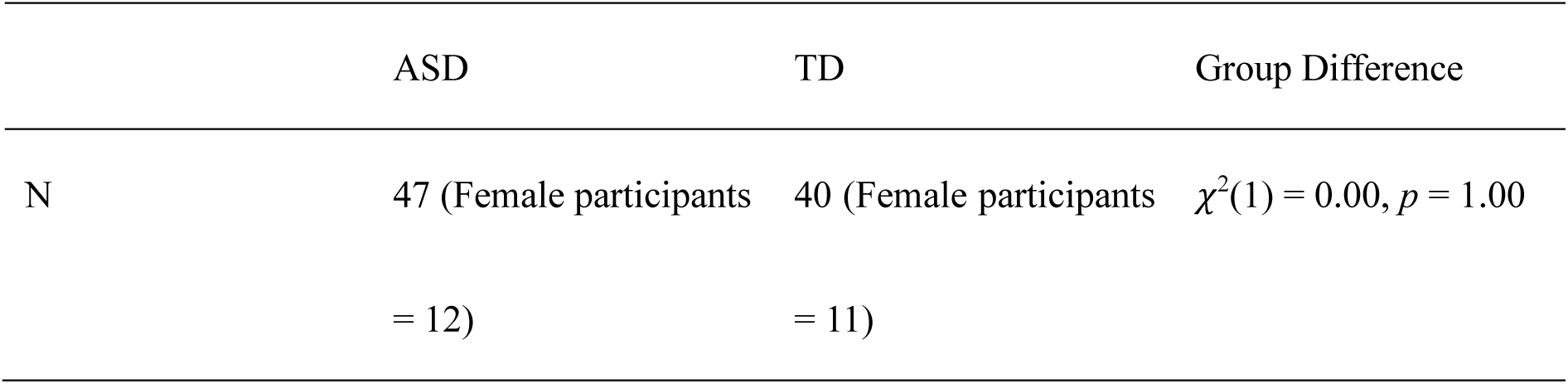
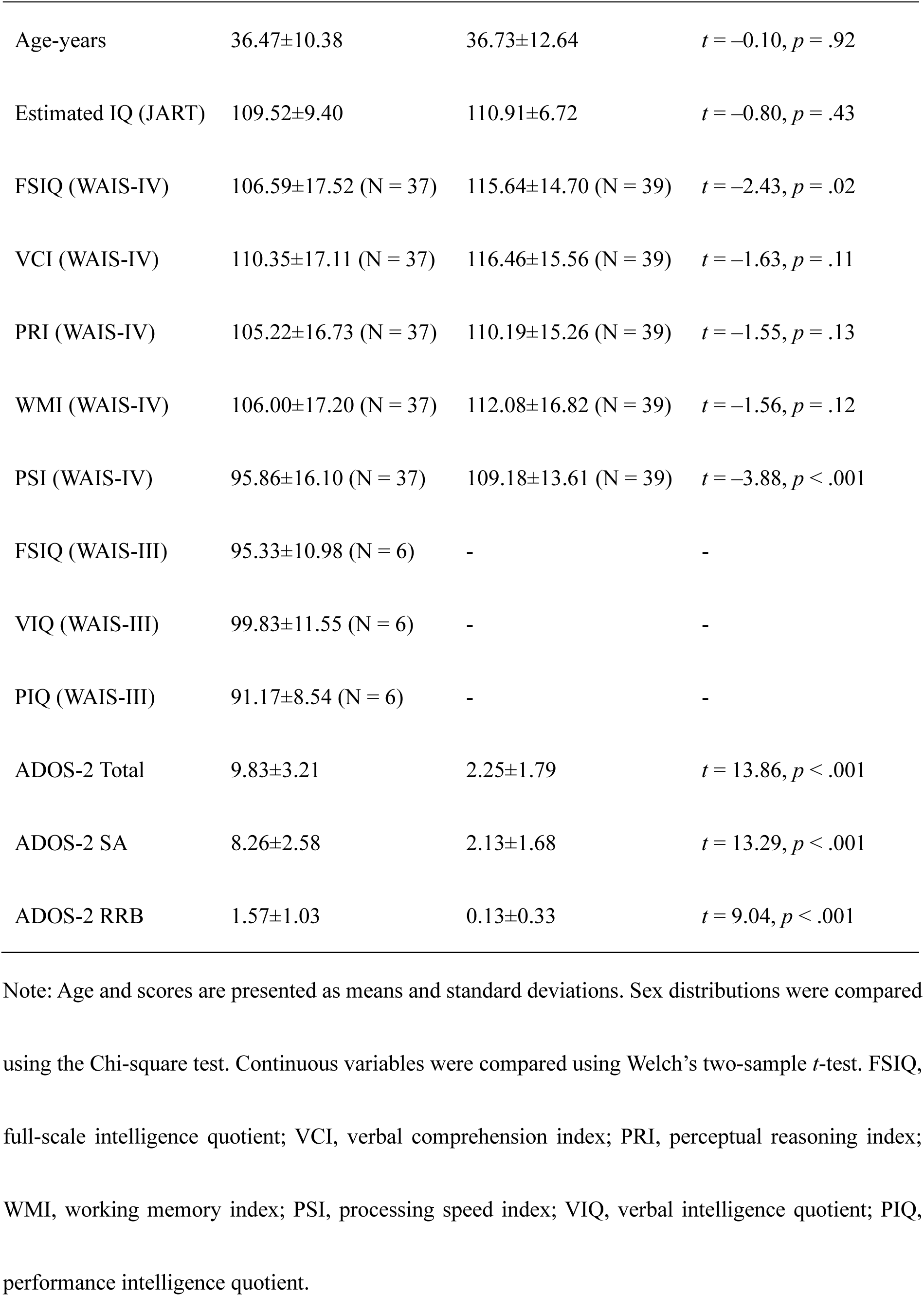
Demographic information and clinical assessment scores.

|  | ASD | TD | Group Difference |
| --- | --- | --- | --- |
| N | 47 (Female participants<br>= 12) | 40 (Female participants<br>= 11) | $\chi^2(1) = 0.00, p = 1.00$ |
| Age-years | 36.47±10.38 | 36.73±12.64 | $t = -0.10, p = .92$ |
| Estimated IQ (JART) | 109.52±9.40 | 110.91±6.72 | $t = -0.80, p = .43$ |
| FSIQ (WAIS-IV) | 106.59±17.52 (N = 37) | 115.64±14.70 (N = 39) | $t = -2.43, p = .02$ |
| VCI (WAIS-IV) | 110.35±17.11 (N = 37) | 116.46±15.56 (N = 39) | $t = -1.63, p = .11$ |
| PRI (WAIS-IV) | 105.22±16.73 (N = 37) | 110.19±15.26 (N = 39) | $t = -1.55, p = .13$ |
| WMI (WAIS-IV) | 106.00±17.20 (N = 37) | 112.08±16.82 (N = 39) | $t = -1.56, p = .12$ |
| PSI (WAIS-IV) | 95.86±16.10 (N = 37) | 109.18±13.61 (N = 39) | $t = -3.88, p < .001$ |
| FSIQ (WAIS-III) | 95.33±10.98 (N = 6) | - | - |
| VIQ (WAIS-III) | 99.83±11.55 (N = 6) | - | - |
| PIQ (WAIS-III) | 91.17±8.54 (N = 6) | - | - |
| ADOS-2 Total | 9.83±3.21 | 2.25±1.79 | $t = 13.86, p < .001$ |
| ADOS-2 SA | 8.26±2.58 | 2.13±1.68 | $t = 13.29, p < .001$ |
| ADOS-2 RRB | 1.57±1.03 | 0.13±0.33 | $t = 9.04, p < .001$ |

Note: Age and scores are presented as means and standard deviations. Sex distributions were compared using the Chi-square test. Continuous variables were compared using Welch's two-sample *t*-test. FSIQ, full-scale intelligence quotient; VCI, verbal comprehension index; PRI, perceptual reasoning index; WMI, working memory index; PSI, processing speed index; VIQ, verbal intelligence quotient; PIQ, performance intelligence quotient.

The sample size was determined based on the number of eligible participants who completed the study during the recruitment period and met the inclusion criteria. No a priori power calculations were performed to determine the analytical sample size. To characterize the sensitivity of the available samples, we conducted a sensitivity power analysis based on a conventional two-group comparison. With 47 participants with ASD and 40 TD participants, a two-sided α of .05 and 80% power, the minimum detectable standardized between-group effect was Cohen’s d = 0.61. This calculation did not use the observed MSE values and should be interpreted as a general benchmark for sample-size sensitivity rather than as a formal power analysis of the covariate-adjusted channel–scale cluster-based permutation procedure.

### Clinical assessment (ADOS-2)

The ADOS-2 was used to quantify ASD-related clinical symptom severity, with particular emphasis on social communication-related symptoms captured by the SA domain. We used revised algorithm scores [48] rather than the original DSM-IV-TR-based algorithm scores. The revised scores were aligned with the DSM-5 criteria. This algorithm evaluates ASD-related behaviors across two domains: SA and Restricted and Repetitive Behaviors (RRB), with the total score across these domains used for diagnostic classification. As outlined in the *Background*, we were particularly interested in the relationship between information-processing characteristics reflected in MSE and social-communication characteristics associated with ASD. Accordingly, the SA score was specified as the primary clinical measure in the analyses that examined the associations between MSE and individual variations in SA characteristics.

### Procedure and experimental stimuli

EEG was recorded during a 7-min resting state with eyes closed (Resting), followed by movie viewing of two video stimuli presented in a fixed order (Table S1): Movie Viewing 1, *Inscapes* [42] (7 min), and Movie Viewing 2, *Partly Cloudy* [43] (5 min, excluding the initial 10 s of production logos). *Inscapes* is a nonnarrative animation developed for neuroimaging research that is composed of slowly evolving abstract shapes and colors [42]. *Partly Cloudy* is a short animated film featuring multiple characters and a coherent character-driven narrative that unfolds through socially and emotionally salient scenes. Previous studies have shown that viewing *Partly Cloudy* reliably engages brain networks implicated in ToM and empathic pain [44, 45]. Silent versions of both videos were used. During the Resting condition, the participants were instructed to close their eyes, relax, avoid focusing on specific thoughts, and remain awake. During both Movie Viewing conditions (Movie Viewing 1 and Movie Viewing 2), participants were instructed to relax and watch the videos. For Movie Viewing 2 (*Partly Cloudy*), participants were additionally instructed to watch the video as they would normally watch a video of interest and attend to its content, because they would later be asked to describe what they had watched verbally^1^. To minimize potential carryover from cognitive engagement during movie viewing to subsequent resting-state MSE, resting-state EEG was recorded before movie viewing, consistent with previous evidence that preceding cognitive or memory engagement can alter subsequent resting-state MSE [53, 54]. The two movie viewing conditions were subsequently presented in a fixed order, with Movie Viewing 1 (*Inscapes*) preceding Movie Viewing 2 (*Partly Cloudy*), to prevent attentional and memory demands associated with the subsequent verbal-description task associated with Movie Viewing 2 from carrying over to Movie Viewing 1.

Participants were seated in front of a 27-inch display (NITRO VG270 bmiix, Acer, Taiwan; refresh rate: 75 Hz). Movie-viewing stimuli were presented on the display, and stimulus presentation, including video playback and synchronization triggers for EEG recording, was controlled using Psychtoolbox [55, 56]. The pixel dimensions of *Inscapes* and *Partly Cloudy*, as determined from the frame size of the original movie files, were 1024 × 800 and 1280 × 720 pixels, respectively.

### EEG data acquisition and preprocessing

EEG signals were acquired using two channel configurations: 64-channel Waveguard™ cap (ANT Neuro, Berlin, Germany) with equidistant scalp positions (ground at the frontal midline anterior to the central channels, reference at 5Z) and a standard 10–10 configuration (ground at AFz/Fpz, reference at Cz/CPz). Although the more widely used 10–10 configuration was preferred for broader applicability, a compatible cap was not available when data collection began. The recordings were initially acquired using the equidistant configuration and transitioned to the 10–10 configuration once the corresponding cap became commercially available. No significant group differences were observed in the proportion of participants using each cap type (equidistant positions: ASD = 33, TD = 27; 10–10 positions: ASD = 14, TD = 13; *χ*^2^[1] = 0.002, *p* = .968). All EEG signals were sampled at 1,000 Hz and band-pass filtered (0.5–70 Hz) throughout data export.

All channels were visually inspected before preprocessing, and channels with unreliable EEG signals were identified and interpolated using spherical spline interpolation. The data were then downsampled to 200 Hz. Line noise at 50 Hz was attenuated using a notch filter, and the data were referenced to the common average. Artifact attenuation was conducted using artifact subspace reconstruction [57], configured to suppress high-amplitude transient artifacts without removing any channel or time segments. Independent component analysis was subsequently applied using the extended Infomax algorithm with dimensionality reduction based on data rank. Components were classified using ICLabel, and those reflecting non-neural activity were removed based on predefined probability thresholds (muscle activity [> 0.70], eye movement [> 0.70], cardiac activity [> 0.80], line noise [> 0.80], and channel noise [> 0.80]). All the EEG data acquired with different channel montages were harmonized using spline-based spatial interpolation. Specifically, the EEG signals were projected from the source montage onto the target montage using spherical spline interpolation of the scalp electric field. The equidistant montage (Fig. S1) was selected as the target space because it provides broader peripheral coverage, thus minimizing extrapolation beyond the spatial extent of the recorded channels, which can introduce distortions in the estimated signals at the outer scalp locations. Finally, current source density [58] transformation was applied using a spherical spline method to enhance the spatial specificity and reduce the effects of volume conduction. Except for channel montage harmonization, EEG preprocessing was conducted using custom MATLAB scripts (MathWorks Inc., Natick, MA, USA) in combination with EEGLAB [59] and FieldTrip [60]. Channel montage interpolation was performed using MNE Python [61, 62].

### MSE analyses

EEG signal complexity was quantified using the MSE [30, 33, 34], which computes sample entropy (SampEn) across scales ranging from fine (or “shorter”) to coarse (or “longer”). SampEn quantifies temporal complexity/irregularity based on the premise that more irregular signals are less likely to preserve pattern similarity as they unfold over time.

Computing the MSE over several minutes of continuous EEG data is computationally demanding; therefore, we analyzed multiple 60-s segments rather than the full recording. Three 60-s EEG segments were extracted from the preprocessed time series (10–70 s, 100–160 s, and 180–240 s). This segmentation was chosen to balance computational tractability with the need to mitigate the influence of temporal fluctuations on neural dynamics during the recordings. The shorter duration of *Partly Cloudy* relative to the other conditions further constrained the segment selection.

For each 60-s time series (*N* = 12,000 time points; sampling rate, 200 Hz), MSE was computed across scale factors (*τ* = 1–30). For an original time series *X* = [*X*_1_, *X*_2_, ⋯, *X_N_*] of length *N*, the coarse-grained time series at scale factor *τ*, denoted 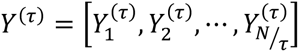 was constructed by averaging consecutive, non-overlapping windows of length *τ*:

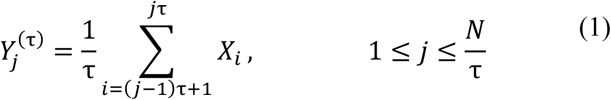

For notational simplicity, *N*/*τ* denotes the number of complete non-overlapping windows at scale factor τ. Thus, *Y*^(1)^ is identical to the original time series, whereas larger *τ* values represent progressively longer temporal scales. At the original sampling rate of 200 Hz, coarse-graining at scale factor τ yields a time series with an effective sampling rate of 200/τ Hz. These effective sampling rates describe the temporal resolution of the coarse-grained time series, and should not be interpreted as conventional EEG frequency bands or direct measures of oscillatory activity at the corresponding frequencies.

SampEn quantifies the negative natural logarithm of the conditional probability that two sequences of *m* consecutive data points that are similar within tolerance *r* remain similar when extended to a length of *m* + 1. Here, *m* denotes the embedding dimension, and *r* is the similarity tolerance. SampEn was computed for each coarse-grained time series, *Y*^(*τ*)^ as follows:

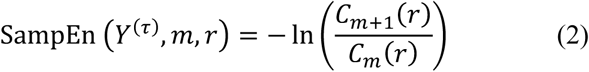

where *C_m_*(*r*) denotes the probability that two sequences of length *m* in the coarse-grained time series *Y*^(*τ*)^ are similar within tolerance *r*. Following prior work examining MSE in neuropsychiatric and neurological disorders [33, 34, 36], the embedding dimension and similarity tolerance were set to *m* = 2 and *r* = 0.2 × the SD of the original time series. These parameter choices are supported by prior studies demonstrating that SampEn provides stable and statistically valid estimates [63, 64]. For each channel and time scale, SampEn values from the three 60-s time series were subsequently averaged and used for statistical analysis to reduce the influence of within-condition temporal fluctuations and obtain a stable estimate of MSE for each condition.

### Statistical analyses

The EEG data were analyzed using custom MATLAB scripts (MathWorks Inc., Natick, MA, USA) in combination with the FieldTrip toolbox [60]. Statistical analyses involving linear mixed-effects (LME) models were conducted using R version 4.4.2 [65], employing the lme4 package [66] for model fitting, the performance package [67] for model diagnostics, the lmerTest package [68] for conventional LME models, the glmmTMB package [69] for models with duration-dependent residual dispersion, the emmeans package [70] for post hoc comparisons, and the effectsize package [71] for the calculation of effect sizes. Since previous studies have reported that EEG MSE varies with age and sex [72–74], all statistical analyses accounted for these variables, as detailed in the respective sections.

### Group difference and effect of experimental condition

To test the effects of Group, Condition (Resting, Movie Viewing 1, and Movie Viewing 2), and their interactions on EEG MSE, we conducted a nonparametric cluster-based permutation analysis across a channel–scale (64 × 30) space.

Before statistical analysis, the effects of age and sex were regressed out of the MSE values at each channel and scale, and the resulting residual MSE values were used for statistical testing. The residual data were arranged as subject × condition × channel × scale arrays, with Group and Condition specified as between- and within-subject factors, respectively. The Group main effect tested whether the residual MSE differed between the ASD and TD participants after collapsing across experimental conditions. The main effect of Condition tested whether the residual MSE differed among the Resting, Movie Viewing 1, and Movie Viewing 2 conditions after collapsing across groups. The Group × Condition interaction tested whether the condition-dependent change in residual MSE differed between ASD and TD participants.

Statistical analyses were conducted using cluster-based nonparametric permutation tests. For each effect, channel–scale points exceeding a cluster-forming threshold of *p* < 0.05 in the upper tail of the *F* distribution were grouped into clusters, and the sum of *F* values within each cluster was used as the cluster statistic. Cluster formation requires suprathreshold effects to extend across at least three neighboring channels. Spatial neighborhoods were defined using the distance-based criterion implemented in FieldTrip. Cluster-level significance was assessed using a Monte Carlo permutation procedure against the permutation distribution of the maximum cluster-wise sum of *F* values obtained from 10,000 permutations. Group labels were permuted for the group effect; condition labels were permuted within participants for the condition effect; and group labels were permuted while preserving each participant’s condition profile for the Group × Condition interaction. Clusters were considered significant at the cluster level *p* < 0.05.

When a significant main effect of the Condition was identified, all three pairwise comparisons among Resting, Movie Viewing 1, and Movie Viewing 2 were conducted using cluster-based non-parametric permutation tests based on t-statistics. The cluster-forming threshold, spatial clustering criteria, number of permutations, maximum cluster statistic procedure, and cluster-level significance threshold were identical to those used in the omnibus *F* statistical analysis. Cluster-level *p* values from all clusters identified across the three pairwise comparisons were jointly corrected using the false discovery rate (FDR) procedure.

Had a significant Group × Condition interaction, we would have tested between-group differences in within-participant condition contrasts for all three pairwise comparisons. These analyses used the same *t*-statistic-based cluster permutation procedure as the follow-up tests for the main effect of Condition. Cluster-level *p* values were corrected across comparisons using the FDR procedure.

For visualization and interpretation, topographical effect-size maps were generated for the omnibus and follow-up analyses. Partial η² was calculated from the observed *F* statistics and plotted for the Group and Condition effects, whereas Hedges’ *g* was calculated from the observed *t* statistics and plotted for the pairwise comparisons.

### Cluster-extent comparison across experimental conditions

We compared the extent of clusters showing significant between-group differences across movie viewing conditions, and examined whether the observed extent difference was robust across cluster-forming thresholds and exceeded that expected under a covariance-matched null model. Following the logic of prior functional magnetic resonance imaging (fMRI) studies comparing the spatial properties of observed thresholded statistical maps with Monte Carlo null maps [75, 76], we assessed the robustness of the spatial-scale extent pattern. Using a mass-univariate approach, we fit a general linear model for each channel–scale feature, including the main effects of Group and Condition, their interactions, and age and sex as covariates. We first used the residuals from this full model to estimate the covariance structure of the unexplained variation. To generate null residuals, we approximated the residual covariance as the Kronecker product of separate channel and scale correlation matrices following covariance modeling approaches developed for EEG and magnetoencephalography data [77]. These maps were rescaled using the residual standard deviation (SD) for each channel–scale feature and added to the null-fitted values. To preserve the within-participant dependence between the two movie viewing conditions, we estimated a single residual correlation between the two conditions and used it to generate correlated condition-specific null maps.

For both the observed data and each null dataset, age- and sex-adjusted group-effect t-maps were computed separately for Movie Viewing 1 and Movie Viewing 2. The threshold sweep included the primary cluster-forming threshold used in the main cluster-based analysis (α = .05) and additionally examined thresholds from α = 0.10-0.01. The α = .05 threshold, matching that used in the primary cluster-based analysis, was treated as the primary threshold for the extent comparison; the additional threshold sweep was used as a robustness analysis to assess the dependence of the result on the choice of cluster-forming threshold rather than as a set of separate confirmatory tests. Suprathreshold clusters were identified using a predefined channel and scale adjacency, and cluster extent was quantified as the number of suprathreshold channel–scale points in the largest cluster. The observed difference in extent, defined as Movie Viewing 2 (*Partly Cloudy*) minus Movie Viewing 1 (*Inscapes*), was compared with the corresponding covariance-matched null distribution.

For visualization, the topographical maps displayed Hedges’ *g*, whereas cluster identification and extent quantification were based on t-maps.

### Sensitivity analyses for channel interpolation and montage harmonization

Channel interpolation and montage harmonization may increase spatial dependence among neighboring channels, thereby influencing the apparent spatial extent of channel-wise cluster-based effects. To assess the robustness of our findings to these procedures, we repeated cluster-based analyses in a restricted dataset comprising only participants whose EEG was recorded using the equidistant scalp position montage (Table S2), excluding all interpolated channels from the analyses. We focused on the equidistant montage because the number of participants recorded with a 10–10 montage was relatively small for a separate statistical analysis.

The number of interpolated channels did not differ between groups (ASD: mean = 0.91, SD = 1.31, range = 0–4; TD: mean = 0.70, SD = 1.07, range = 0–4; Welch’s *t* = 0.67, *p* = .506). Interpolated channels were generally sparse across the montage in both groups (Table S3), with the highest frequencies of missingness observed at the posterior scalp locations 11 L (ASD, 3/33; TD, 6/27) and 11R (ASD, 4/33; TD, 5/27).

### Leave-one-subject-out identification of group-difference clusters and their associations with ADOS-2 scores

We examined whether MSE within the channel–scale cluster showing significant between-group differences was associated with individual variations in SA characteristics, as indexed by ADOS-2 SA scores.

Since extracting neural values from a cluster defined using the full sample would use each participant’s data for both cluster selection and neural value extraction, the independence of these two steps would not be preserved [78]. To mitigate circularity, we used a leave-one-subject-out (LOSO) procedure adapted from non-circular feature selection approaches in fMRI region-of-interest studies [79]. For each condition, the MSE at each channel–scale feature was calculated separately for three 60-s EEG segments extracted from the preprocessed time series (10–70, 100–160, and 180–240 s) and then averaged for each participant. The condition-specific MSE values were averaged across the three conditions. In each LOSO fold, one participant was excluded, and the resulting participant-level means from the remaining sample were entered into a general linear model that included group, age, and sex. Therefore, the Group *t*-statistic tested the age- and sex-adjusted between-group differences averaged across conditions, corresponding to the group main effect in the omnibus analysis. Clusters were identified from these *t*-statistics using a two-sided cluster-based permutation procedure with 10,000 permutations, a cluster-forming threshold of *p* < 0.05, and a cluster-level significance threshold of *p* < 0.05. The resulting significant cluster mask was applied to the held-out participant, for whom SampEn was averaged across all channels and scales within the cluster. Repeating this procedure for every participant yielded participant-level cluster-averaged SampEn estimates for subsequent analyses, with mask selection independent of each participant’s held-out data.

For each participant and condition, the MSE values were averaged across all channel–scale points within the LOSO-derived group-difference cluster, yielding a single cluster-averaged MSE value for each condition. We then fitted LME models to examine whether MSE within the LOSO-derived cluster was associated with individual variation in ADOS-2 SA scores, and whether these associations differed across diagnostic groups or experimental conditions.

Binary predictors, including Group and sex, were effect-coded as −0.5 and 0.5. Condition was represented by sum-to-zero contrasts so that the main effect of the ADOS-2 SA score reflected its association with MSE averaged across the three conditions. The cluster-averaged MSE and age were standardized to have a mean of zero. ADOS-2 SA scores were centered within each diagnostic group to remove between-group differences in score levels and isolate variations relative to each group-specific mean, following the general principles of group-mean centering [80]. The centered scores were then divided by the pooled within-group SD, calculated by weighting the group-specific variances by their respective degrees of freedom [81, 82].

The fixed-effects structure included ADOS-2 SA, Group (ASD or TD), Condition (Resting, Movie Viewing 1, or Movie Viewing 2), ADOS-2 SA × Group, and ADOS-2 SA × Condition. Age and sex were included as nuisance covariates, and random intercepts were specified for each participant. The LME models satisfied the commonly used diagnostic criteria, including low variance inflation factors (VIFs < 5), successful convergence, and no evidence of singular fit.

### Event-related MSE changes during socially relevant and temporally matched non-social scenes

We conducted hypothesis-driven follow-up analyses within channels and at longer scales, which contributed to the overall group effect. These analyses aimed to determine whether the group-difference pattern in MSE differed during socially relevant events and whether such event-related changes were associated with individual variations in ADOS-2 SA scores. Since the preceding analysis was based on a 60-s time series and did not isolate specific socially relevant events, we examined event-related MSE changes during empathic pain and ToM events in *Partly Cloudy*, which have previously been shown to engage corresponding social brain networks. These changes were compared with those observed during temporally matched non-social windows in *Inscapes*.

To avoid circularity, the event-level analysis used the same LOSO-derived group-difference cluster defined in the preceding section.

Socially relevant events in the *Partly Cloudy* condition were defined using time windows previously shown to reliably elicit activity in the brain networks implicated in empathic pain and ToM [45]. Specifically, a previous fMRI study used a data-driven reverse-correlation procedure to identify time windows independently in two adult samples. We used windows identified in the first sample that showed at least a partial temporal overlap with those independently identified in the second sample. As described in the Procedure and Experimental Stimuli section, the corresponding time windows were shifted to account for this temporal offset, because the first 10 s of the movie were removed. Additionally, we verified the selected windows by comparing the event descriptions reported by Richardson et al. (2018) [45] with the corresponding scenes in an edited movie. This yielded seven ToM events, each lasting 4–16 s (mean duration, 9.7 s), and nine empathic-pain events, each lasting 4–18 seconds (mean duration, 8.2 s). The details of the selected windows are presented in Table S7. The first 20 s of the movie, which did not overlap with any target event, served as the control window [83].

Event-related MSE was extracted separately for each participant as follows. MSE was calculated for each channel using the EEG time series corresponding to each target event and control window. To isolate event-related changes associated with empathic pain and ToM processing, MSE during the control window was subtracted from MSE during each target event [83]. For each resulting difference, the values were averaged across the channels and scales included in the participant-specific LOSO-derived clusters. The same procedure was applied to the temporally matched windows in *Inscapes*, thereby controlling for the potential effects of temporal progression on MSE. Therefore, each participant contributed 32 cluster-averaged MSE change scores: nine empathic pain and seven ToM events for each of the two stimulus conditions: *Partly Cloudy* (social) and *Inscapes* (non-social).

LME models were used to test event-related MSE changes, diagnostic group differences, and associations with individual variations in ADOS-2 SA scores. Binary predictors were effect-coded as −0.5 and 0.5. The cluster-averaged MSE change, age, and time window duration were standardized to a mean of zero and an SD of one. ADOS-2 SA scores were centered within the diagnostic group and subsequently scaled using the pooled within-group SD. The cluster-averaged MSE change was used as the dependent variable. The fixed-effects structure included the main effects of Group (ASD or TD), Stimulus (nonsocial or social), Event (empathic pain or ToM), and ADOS-2 SA score; all interactions among Group, Stimulus, and Event; and all interactions among ADOS-2 SA score, Stimulus, and Event. Time window duration, participant age, and sex were included as nuisance covariates. Random intercepts were specified for each participant and time window. The model converged successfully, showed no evidence of a singular fit, and exhibited low multicollinearity (all VIFs < 5). For the Group × Stimulus interaction and follow-up Stimulus contrasts, standardized effect sizes were expressed as partial correlation coefficients (partial *r*), approximated from the corresponding *t-*statistics and denominator degrees of freedom.

It is important to note that a reliable MSE estimation requires a sufficient number of continuous data points, particularly at longer scales, where coarse-graining progressively reduces the effective length of the time series. Although no universal minimum applies [84], previous studies have used a time series with as few as 50 points per scale [85], whereas heuristic recommendations have suggested that at least 100 and preferably 900 points may be needed for a stable estimation of the underlying conditional probabilities [86]. In the present dataset, the shortest segment was 4 s (800 samples at 200 Hz), yielding 26 coarse-grained data points at the longest scale (τ = 30). To assess the stability of MSE estimates for events of the same type, we performed supplementary analyses to examine whether MSE values and the relative ranking of participants were preserved as progressively shorter segments were extracted from the relatively long event-related time windows (Supplementary Data 3).

## Results

### Group difference and effect of experimental condition

A significant cluster for the main effect of Group was identified (cluster-level *p* = .025), indicating lower SampEn at longer scales (τ = 16–30, coarse-grained time series with effective sampling rates of 12.50–6.67 Hz) in the ASD group than in the TD group (Fig. 1). This cluster included channels showing effects ranging from small to large (partial η^2^ = 0.045 to 0.133). Fig. 1B shows the mean MSE curves across the channels included in this cluster, and Fig. S2 shows the distribution of participant-level cluster-averaged MSE values.

**Fig. 1.**
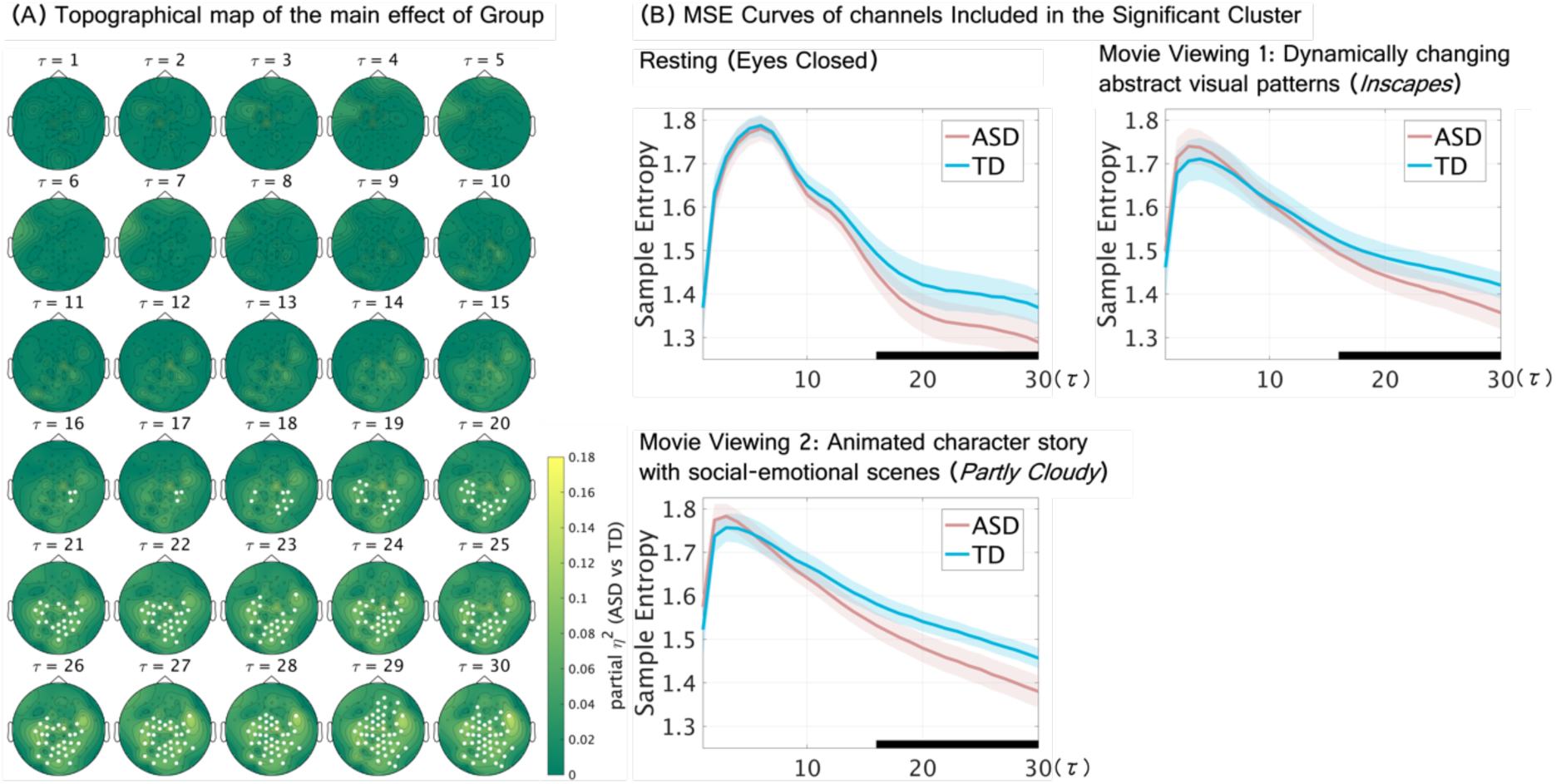
Group (ASD/TD) differences in EEG multiscale entropy (MSE). (A) Topographical maps showing channel–scale partial *η²* effect-size maps for the Group effect, derived from *F*-statistics computed from age- and sex-adjusted SampEn values. Each topographical map represents the between-group differences in SampEn, computed from time series obtained by coarse–graining the original signal to 200/τ Hz. τ denotes the scale factor. A single significant cluster showing a Group effect was identified, and channels belonging to this cluster are highlighted in white. Statistical significance was assessed using one–sided cluster-based permutation tests (10,000 permutations; cluster-forming threshold *p* < 0.05; cluster–level *p* < 0.05). (B) Separate line plots for each condition show mean SampEn averaged across all channels included in the significant cluster at one or more scales (ASD: pink, TD: sky blue), with shaded bands indicating the 95% confidence intervals. In each plot, a black bar along the x-axis indicates the range of time scales spanned by the significant cluster.

A significant main effect of Condition was identified in a cluster (cluster-level *p* < .001) spanning shorter-to-longer scales (*τ* = 4–30, coarse-grained time series with effective sampling rates of 50–6.67 Hz), indicating that MSE varied across conditions irrespective of Group (Fig. S3). Partial η² values indicated small-to-large Condition effects across the cluster (partial η^2^ = 0.035 to 0.436). Post hoc cluster-based permutation tests identified significant clusters for all three pairwise condition contrasts (FDR-corrected cluster-level *ps* ≤ .01). Topographical maps showing the channels included in the significant clusters for each contrast together with the corresponding effect sizes for each channel are shown in Fig. S4A–C. Since the significant clusters showed a widespread scalp distribution, the mean MSE values averaged across all channels are plotted for each condition in Fig. S4D.

Resting versus Movie Viewing 1 and Resting versus Movie Viewing 2 showed broadly similar patterns, with three significant clusters identified in each contrast: MSE was lower during Resting at short scales (vs Movie Viewing 1: τ = 1–2; corresponding to coarse-grained time series with effective sampling rates of 200–100 Hz; FDR-corrected cluster-level p = .009; Hedges’ g = –0.938 to –0.226; vs Movie Viewing 2: τ = 1–3; corresponding to coarse-grained time series with effective sampling rates of 200–66.67 Hz; FDR-corrected cluster-level p = .002; Hedges’ g = –1.269 to –0.212) and longer scales (vs Movie Viewing 1: τ = 15–30; corresponding to coarse-grained time series with effective sampling rates of 13.33–6.67 Hz; FDR-corrected cluster-level p < .001; Hedges’ g = –0.863 to –0.214; vs Movie Viewing 2: τ = 12–30; corresponding to coarse-grained time series with effective sampling rates of 16.67–6.67 Hz; FDR-corrected cluster-level p < .001; Hedges’ g = –1.238 to –0.212), whereas MSE was higher during Resting at short-to-middle scales (vs Movie Viewing 1: τ = 5–14; corresponding to coarse-grained time series with effective sampling rates of 40–14.29 Hz; FDR-corrected cluster-level p = .002; Hedges’ g = 0.212 to 0.812; vs Movie Viewing 2: τ = 5–9; corresponding to coarse-grained time series with effective sampling rates of 40–22.22 Hz; FDR-corrected cluster-level p = .006; Hedges’ g = 0.216 to 0.720). The comparison between the two Movie Viewing conditions identified a significant cluster spanning the full range of scales, indicating lower MSE during Movie Viewing 1 than during Movie Viewing 2 (τ = 1–30, corresponding to coarse-grained time series with effective sampling rates of 200– 6.67 Hz; FDR-corrected cluster-level p < .001; Hedges’ g = –0.695 to –0.212).

No significant clusters were observed for the Group × Condition interaction.

The Group and Condition effects were broadly reproduced when analyses were restricted to participants recorded with the equidistant scalp-position montage and interpolated channels were excluded (Figs. S5–S7).

### Cluster-extent comparison across experimental conditions (Movie Viewing 1 vs Movie Viewing 2)

Fig. 2A presents the ASD–TD Hedges’ *g* maps separately for the Movie Viewing 1 and Movie Viewing 2 conditions, with suprathreshold clusters identified using a cluster-forming threshold of α = .05.

**Fig. 2.**
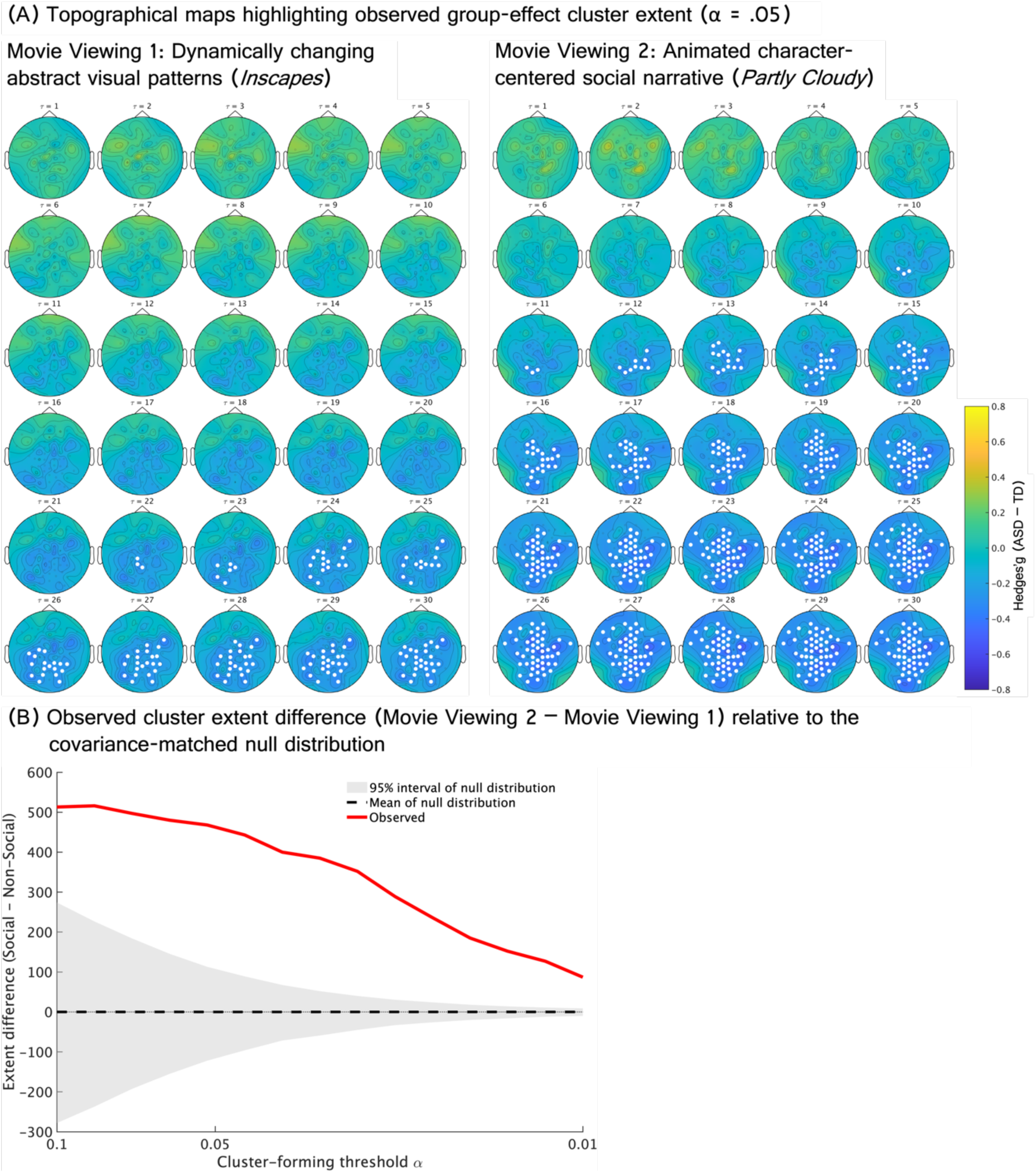
Differences in between-group cluster extent between Movie-Viewing conditions (Movie Viewing 1 vs Movie Viewing 2). (A) Topographical channel–scale maps show Hedges’ g (ASD–TD) maps in each condition. Each topographical map represents the Hedges’ g of between-group differences in SampEn, computed from time series obtained by coarse–graining the original signal to 200/*τ* Hz. *τ* denotes the scale factor. Channels belonging to the observed suprathreshold cluster are highlighted in white. (B) Observed cluster extent differences are shown across cluster-forming thresholds from α = 0.10 to α = 0.01. Cluster extent was quantified as the number of suprathreshold channel–scale points in the largest cluster. The covariance-matched null expectation was obtained from Monte Carlo Gaussian random fields preserving the empirical channel–scale covariance structure.

At the primary cluster-forming threshold (α = .05), the largest suprathreshold cluster was 453 channel–scale points more extensive during Movie Viewing 2 than during Movie Viewing 1 (two-sided Monte Carlo *p* = .004). In the robustness analysis, the extent difference remained positive across cluster-forming thresholds from α = .10 to .01, ranging from 87 to 517 channel–scale points (Fig. 2B). The observed differences also remained outside the corresponding covariance-matched Monte Carlo null distributions across the threshold sweep (two-sided Monte Carlo *ps* ≤ .014).

These between-condition differences in cluster extent were broadly reproduced when analyses were restricted to participants recorded with the equidistant scalp-position montage and interpolated channels were excluded (Fig. S8).

### Associations of LOSO-derived group-difference cluster MSE with ADOS-2 SA scores

The LOSO-derived probability map showed a substantial spatial and scale-wise overlap with the group-difference cluster identified in the full-sample analysis (Fig. 3A).

**Fig. 3.**
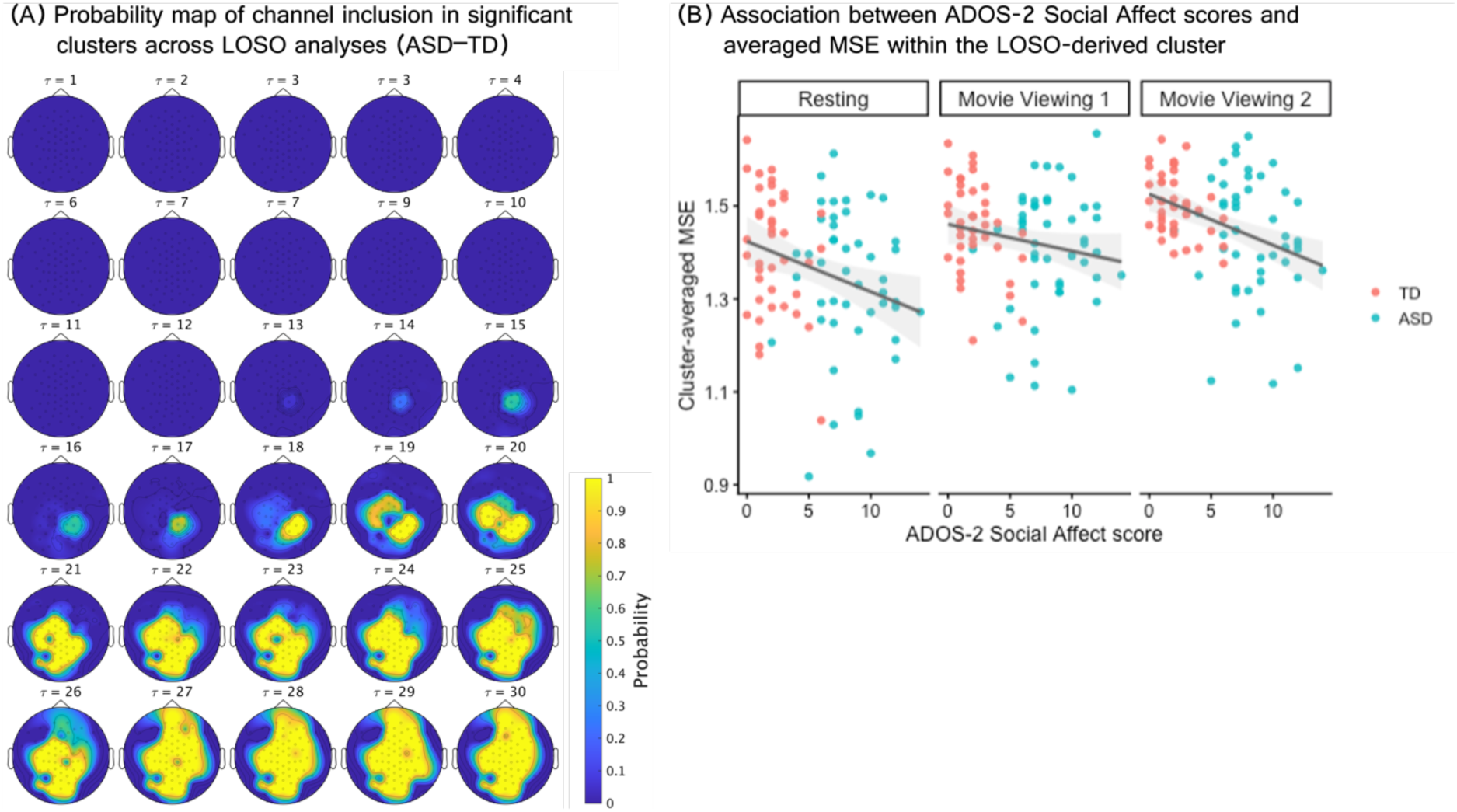
Association between the group-difference pattern in MSE and ADOS-2 Social Affect scores. (A) Topographical maps show the probability that each channel was included in the significant (ASD– TD) cluster across the leave-one-subject-out (LOSO) iterations. Each topographical map represents the inclusion probabilities computed from time series obtained by coarse–graining the original signal to 200/*τ* Hz. *τ* denotes the scale factor. (B) Scatter plot showing the relationship between ADOS-2 Social Affect scores and mean SampEn across channels and scales within the LOSO-derived cluster. Lines indicate descriptive linear fits with 95% CIs. Condition-specific simple-slope analyses indicated significant negative associations between ADOS-2 Social Affect scores and cluster-averaged SampEn during Resting and Movie Viewing 2 (FDR-corrected *ps* = .035). In contrast, the association was not significant during Movie Viewing 1 (FDR-corrected *p* = .339).

LME revealed a significant negative main effect of ADOS-2 SA on cluster-averaged MSE, indicating that higher ADOS-2 SA scores were associated with lower MSE across the experimental conditions and diagnostic groups (Estimate = –0.037, 95% confidence interval [CI] = [–0.071, –0.003], *p* = .032). Neither the ADOS-2 SA × Group interaction nor the ADOS-2 SA × Condition interaction (*ps* ≥ .057) was statistically significant. To characterize the condition-specific pattern of association, we conducted simple exploratory slope analyses for each experimental condition. These analyses indicated significantly negative associations between ADOS-2 SA scores and MSE in the Resting condition (*β*= –0.255, 95% CI = [–0.475, –0.035], FDR-corrected *p* = .035) and during Movie Viewing 2 (*Partly Cloudy*; *β*= –0.272, 95% CI = [–0.492, –0.052], FDR-corrected *p* = .035). The estimated slope was also negative during Movie Viewing 1 (*Inscapes*), although it was not statistically significant (*β*= –0.107, 95% CI = [–0.327, 0.113], FDR-corrected *p* = .339). It should be noted, however, that the omnibus ADOS-2 SA × Condition interaction was not significant. Fig. 3B shows the associations between the ADOS-2 SA scores and MSE in each experimental condition. The complete results from the LME model, including fixed-effect estimates and statistical tests for the nuisance covariates age and sex, are reported in Table S4. Corresponding analyses using the ADOS-2 RRB and total scores are also reported in Tables S5 and S6, respectively; neither score showed a significant association with MSE.

### Event-related MSE changes during socially relevant and temporally matched non-social scenes

The LME model revealed significant Group × Stimulus and Stimulus × ADOS-2 SA score interactions (Group × Stimulus: Estimate = –0.166, 95% CI = [–0.289, –0.046], *p* = .008; Stimulus × ADOS-2 SA score: Estimate = –0.093, 95% CI = [–0.174, –0.004], *p* = .026). The Group × Stimulus interaction indicated that the within-participant social-minus-non-social difference in the cluster-averaged MSE change was smaller in the ASD group than in the TD group (*partial r* = –0.051, 95% CI = [–0.089, –0.013], *p* = .008; Fig. 4). Follow-up contrasts showed that the longer-scale cluster-averaged MSE change was significantly higher during social stimuli than during non-social stimuli in the TD group (*partial r* = 0.092, 95% CI = [0.054, 0.129], FDR-corrected *p* < .001). In the ASD group, the estimated difference did not reach statistical significance (*partial r* = 0.023, 95% CI = [–0.015, 0.061], FDR-corrected *p* = .237). Although the standardized effect sizes were small, these findings indicate that social processing-related changes in longer-scale MSE differed between the diagnostic groups. The post hoc analysis of the Stimulus × ADOS-2 SA score interaction indicated that the condition-specific slopes relating ADOS-2 SA scores to cluster-averaged MSE changes were not statistically significant in either condition (|*partial rs*| < 0.104, FDR-corrected ps > .641). However, the difference between the slopes for the Social and Non-Social stimuli was statistically significant *(partial r* = 0.043, 95% CI = [0.005, 0.081], p = .026). No effects involving Event (corresponding to empathic pain and ToM events under *Partly Cloudy*) were statistically significant. The full results of the LME model, including the nuisance covariates, are presented in Table S8.

**Fig. 4.**
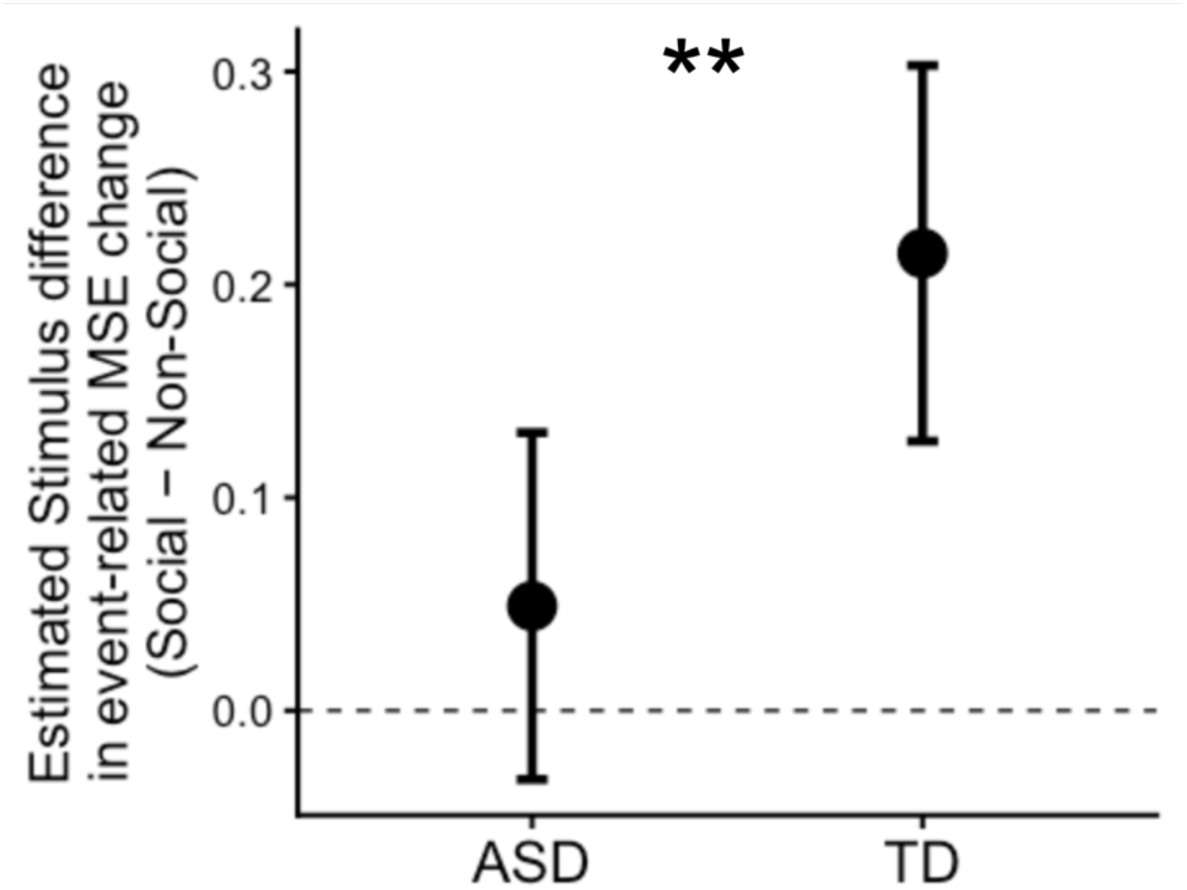
Group × Stimulus interaction for the cluster-averaged event-related MSE change. Black dots indicate the model-estimated Social–Non-Social difference in cluster-averaged event-related MSE change, comparing socially relevant events in *Partly Cloudy* with temporally matched non-social windows in *Inscapes*, averaged across ToM and empathic-pain events. Error bars indicate 95% CIs. \*\**p* < .01.

Supplementary analyses examining the stability of MSE estimates showed that even short excerpts generally preserved relative between-participant differences, whereas the 4-s estimates showed relatively higher estimation error and variability, and somewhat reduced numerical estimability (Figs. S9–12). Therefore, we conducted two complementary sensitivity analyses: one retaining all windows while modeling residual variance as a function of excerpt duration, and another applying the same LME model as in the primary analysis after excluding the 4-s windows to address their comparatively lower numerical estimability. In both models, the Group × Stimulus interaction remained significant, with effect directions consistent with those observed in the primary analysis (Tables S9 and S10).

## Discussion

In this study, we found that longer-scale MSE was lower in adults with ASD than in TD adults and was negatively associated with ADOS-2 SA scores; the ASD–TD differences spanned a broader spatial-by-scale domain during viewing of a socioemotional animated narrative than during viewing of abstract visual patterns, and the increase in MSE during social relative to non-social events was smaller in ASD than in TD participants. These findings were obtained after accounting for age and sex, both of which have been previously reported to influence MSE [72–74].

The reduction in long-scale MSE observed in adults with ASD may represent a broadly expressed neurophysiological feature of ASD. Although previous studies have reported reduced MSE in ASD, particularly at longer scales, most have examined infants or children, and recording conditions have varied considerably across studies, including during eyes-open or eyes-closed rest and cognitive-task performance [5, 37–41]. The present study replicated and extended previous evidence of reduced longer-scale MSE in adults with ASD by demonstrating descriptively consistent ASD-TD differences across all three conditions. These convergent findings suggest that reduced longer-scale MSE in ASD is not confined to a particular developmental stage or recording state but may reflect a comparatively stable feature of the condition. Given that alterations in conventional EEG measures in ASD often vary with developmental stage and recording context [19, 20], measures of EEG irregularity and complexity, and more broadly, the temporal dynamics of neural activity, may provide candidate neurophysiological features of ASD.

The spatial and scale-wise patterns of the ASD–TD difference may vary across movie viewing conditions, although there was no evidence of a Group × Condition interaction in effect magnitude. The cluster showing ASD-TD differences was more spatially and scale-wise extensive during the viewing of the socio-emotional narrative (*Partly Cloudy*) than during the viewing of abstract visual patterns (*Inscapes*). A consistent pattern was observed at the primary cluster-forming threshold, which remained directionally consistent throughout the threshold-sweep robustness analysis. These findings suggest that group differences vary across experimental contexts, although the physiological mechanisms underlying this variation remain unknown.

The group-difference and cluster-extent findings were broadly reproduced in sensitivity analyses restricted to a single montage and excluding the interpolated channels. These findings suggest that potential spatial expansion bias arising from channel interpolation and montage harmonization did not substantially influence the observed results.

Using participant-specific LOSO-derived masks, which excluded each participant’s data from the cluster definition to minimize circularity, the MSE within the longer scales and channels contributing to the ASD–TD difference was lower in individuals with higher ADOS-2 SA scores. This association was observed after removing between-group differences in ADOS-2 SA scores, indicating a dimensional relationship between neural complexity and social-affect characteristics within diagnostic groups, with no statistical evidence that the slope differed between ASD and TD. Although this analysis does not directly test a continuum spanning ASD and TD, the observed pattern may be compatible with the view that ASD-related traits are continuously distributed in the general population [87–89]. In the supplementary analyses, we applied the same analyses to ADOS-2 RRB and total scores. Neither score showed a significant association with the longer-scale MSE. Thus, the present study provides no evidence of an association between long-scale MSE and RRB, the other core symptom domain of ASD. Since SA and RRB were analyzed separately and may co-vary, the present findings do not establish the domain specificity of the MSE association.

The event-related analyses further suggest that longer-scale MSE may capture differences in social information processing. We compared event-related changes in longer-scale MSE, defined as MSE during each target segment minus MSE during an initial control segment of the same movie, between empathic-pain and ToM scenes in *Partly Cloudy* and non-social segments occurring at the same movie-relative time points in *Inscapes*. The Social-minus-Non-Social difference in MSE change was significantly smaller in ASD participants than in TD participants. Follow-up analyses showed no significant difference in ASD participants, whereas TD participants showed significantly greater MSE changes during social events than during non-social events. Social communication requires the integration of multiple continuously evolving signals and the flexible adjustment of behavior in response to others [28, 29]. Such computational demands may recruit a broader repertoire of temporal dynamics, thereby increasing neural complexity, as was clearly observed in TD participants. Therefore, difficulties in social communication in ASD may be related, at least in part, to a reduced capacity to flexibly reconfigure neural dynamics when processing complex and rapidly changing social information. This interpretation highlights the importance of examining how neural dynamics relate to information processing in ASD in the context of reduced spontaneous orientation towards others’ intentions and emotions [90–93] and atypical empathic processing [94–100]. Longer-scale MSE did not distinguish between empathic pain and ToM events. Future studies combining MSE with complementary measures of rapid neural dynamics [101] may clarify whether distinct social processes involve different patterns of temporal reconfiguration. Nevertheless, the interpretation of the present event-related findings requires careful consideration of potential confounding factors. *Partly Cloudy* and *Inscapes* differed not only in social content but also in narrative structure, task instructions, and native pixel dimensions, among other stimulus properties. Defining event-related MSE changes relative to a control segment within each movie may have reduced the influence of stable between-movie differences. However, it cannot fully isolate social information processing from other stimulus characteristics. Future studies using scrambled, temporally reordered, or reverse-played stimuli could help separate the effects of social processing from other stimulus-related confounders.

Based on these findings, we considered the cognitive and physiological mechanisms that may contribute to the observed MSE patterns in ASD. In contrast to the reduced longer-scale MSE identified in ASD, previous studies have reported increased longer-scale MSE in schizophrenia [34] and Alzheimer’s disease [33]. These divergent MSE patterns may reflect differences in the underlying synaptic or circuit-level organization. ASD has been associated with increased dendritic spine density, potentially related to incomplete synaptic pruning, whereas schizophrenia has been associated with excessive spine pruning, and Alzheimer’s disease with progressive synaptic loss [102]. In principle, these distinct patterns of synaptic alterations could contribute to differences in large-scale neural dynamics. Within this framework, reduced longer-scale MSE in individuals with ASD may reflect less flexible neural dynamics and may relate to difficulties in integrating multimodal and context-dependent social information. In contrast, increased longer-scale MSE in schizophrenia [34] may reflect dysregulated neural dynamics characterized by insufficient constraints, rather than greater adaptive flexibility, a pattern that could be relevant to disorganized thought and speech [1, 2]. Notably, scalp EEG MSE cannot identify the underlying cellular mechanisms, and these mechanistic interpretations remain speculative. The plausibility of this hypothesis should be evaluated in future studies to clarify how scale-specific MSE measured from scalp EEG relates to structural features of the brain, including synaptic density. The Condition effects observed in the present study partially agree with previous EEG-MSE findings. Rest-to-task transitions have been reported to reverse direction across temporal scales, with greater MSE at rest over shorter or intermediate scales but greater task-related MSE at longer scales [103]; another study reported greater task-related MSE at fine scales but greater resting-state MSE at intermediate scales [104]. Greater MSE has also been observed during social face processing than during non-social object processing [5]. This study extends previous work by demonstrating complementary scale-dependent MSE patterns within the same participants across resting, nonsocial movie viewing, and socioemotional movie viewing conditions. The broadly similar condition-dependent profiles in the ASD and TD groups, together with the absence of a significant Group × Condition interaction, suggest that these effects reflect general state-dependent changes in MSE. Collectively, these findings indicate that experimental context does not uniformly increase or decrease MSE, but instead reshapes its multiscale profile.

### Limitations

This study had several limitations. First, the experimental conditions were presented in a fixed order, with the resting condition followed by *Inscapes* and then *Partly Cloudy*. Therefore, the condition effects cannot be fully separated from potential order-related influences, including habituation, fatigue, and carryover. In addition, the two movie stimuli differed not only in social content but also in narrative structure, task instructions, and visual properties. Although event-related MSE changes were defined relative to within-movie control windows and temporally matched across movies, these procedures could not fully isolate social information processing from other stimulus-related differences.

Second, the event-level MSE was estimated from relatively short EEG segments (4–18 s). Although sensitivity analyses accounting for duration-dependent residual dispersion and excluding the 4-s windows supported the robustness of the Group × Stimulus interaction, MSE estimates from the shortest windows showed greater estimation error and variability. This limitation is particularly relevant at coarser scales, where coarse graining further reduces the number of available data points. Therefore, the event-level findings should be interpreted primarily in terms of the relative social–nonsocial contrast rather than as indicating comparable precision of absolute MSE estimates across window durations.

Third, the modest sample size limited the precision of the effect estimates, particularly for small effects and higher-order interactions. Independent replication cohorts were not available for this study. Although the Group and Condition findings and event-level Group × Stimulus interactions were supported by complementary sensitivity analyses, these analyses assessed robustness within the present sample and did not constitute an independent replication. Moreover, some secondary associations involving ADOS-2 scores showed relatively modest statistical evidence and should, therefore, be interpreted in the context of their effect sizes and CIs. To our knowledge, no external dataset was available that combined a sufficiently comparable EEG protocol, the same within-participant resting-state and naturalistic movie-viewing conditions, and the relevant clinical measures. Although the principal findings were robust across leave-one-subject-out and sensitivity analyses, confirmation in an independent cohort is needed. Confirmation in larger, independent adult cohorts is important.

Fourth, female participants were underrepresented in both groups, which may limit generalizability to more sex-balanced populations, despite statistical adjustment for sex. In addition, 14 participants with ASD were taking psychotropic or neurological medications; therefore, potential medication-related influences on MSE could not be excluded.

Fifth, although the LOSO procedure ensured that each participant’s data were excluded when defining the channel–scale mask used to extract the participant’s MSE, it remained an internal feature selection procedure based on largely overlapping training samples. Independent datasets will be required to establish the reproducibility and generalizability of the identified channel–scale patterns.

More broadly, the relationship between MSE and the functional aspects of the ASD phenotype is not fully understood. In the present study, the association between longer-scale MSE and scores related to reciprocal social communication, a core domain of ASD, was statistically significant. We also examined RRB, another core symptom domain of ASD. No statistically significant association was found between longer-scale MSE and RRB scores. Clarifying the relationship between MSE and RRB may require larger samples and experimental paradigms that directly assess neural activity during RRB-relevant states, such as sensory seeking, insistence on sameness, and stereotyped behavioral tendencies.

ASD is a complex and heterogeneous neurodevelopmental condition characterized by variability in trait severity, with substantial diversity in co-occurring conditions, including attention-deficit/hyperactivity disorder, epilepsy, anxiety, depression, and cognitive profiles [1, 2, 105, 106]. Examining how these multidimensional aspects of the ASD phenotype are related to MSE can provide important insights into the neurophysiological basis of ASD heterogeneity.

Finally, the present study did not directly assess synaptic or circuit-level mechanisms. MSE is an indirect system-level measure of signal complexity, and the present data cannot be used to determine whether the observed differences reflect synaptic pruning, synaptic density, or other circuit properties. Future studies that integrate EEG complexity measures with more direct measures of neurobiological structure and function are needed to evaluate these possibilities.

## Conclusions

We examined MSE in adults with ASD and TD during eyes-closed rest and two movie-viewing conditions. Adults with ASD showed reduced MSE at longer scales across resting and naturalistic movie-viewing conditions. Across the diagnostic groups, lower longer-scale MSE was associated with higher ADOS-2 SA scores. Event-level analyses further showed a smaller social–nonsocial difference in longer-scale MSE changes in ASD participants than in TD participants. These findings identify longer-scale EEG complexity as a candidate neurophysiological feature associated with social-affect characteristics in ASD, although confirmation in larger, independent cohorts is needed.

## Abbreviations

ASD: autism spectrum disorder
EEG: electroencephalography
MSE: multiscale entropy
SampEn: sample entropy
TD: typically developing
ToM: theory-of-mind
ADOS-2: Autism Diagnostic Observation Schedule, Second Edition
SA: Social Affect
RRB: Restricted and Repetitive Behaviors
DSM-5: the Diagnostic and Statistical Manual of Mental Disorders, Fifth Edition
DSM-IV-TR: the Diagnostic and Statistical Manual of Mental Disorders, Fourth Edition, Text Revision
WAIS-III: the Wechsler Adult Intelligence Scale–Third Edition
WAIS-IV: the Wechsler Adult Intelligence Scale– Fourth Edition
IQ: Intelligence Quotient
JART: the Japanese version of the National Adult Reading Test
SD: standard deviation
FSIQ: Full Scale Intelligence Quotient
VCI: Verbal Comprehension Index
PRI: Perceptual Reasoning Index
WMI: Working Memory Index
PSI: Processing Speed Index
VIQ: Verbal Intelligence Quotient
PIQ: Performance Intelligence Quotient
LME: linear mixed-effects
FDR: the false discovery rate
LOSO: leave-one-subject-out
fMRI: functional magnetic resonance imaging

## Declarations

### Ethics approval and consent to participate

The Institutional Review Board of Showa Medical University (approval no. 2023-181-A) and the Institutional Review Board of Showa Medical University Karasuyama Hospital (approval no. B-2021-007) approved all procedures conducted in this study. Written informed consent was obtained from all the participants after fully explaining the purpose of this study. The authors assert that all procedures contributing to this work comply with the ethical standards of the relevant national and institutional committees on human experimentation and with the Helsinki Declaration.

### Consent for publication

All participants provided consent for publication.

### Availability of data and materials

The datasets used and/or analyzed during the current study are available from the corresponding author on reasonable request.

### Authors’ contributions

TN (Conceptualization, Investigation, Data curation, Formal analysis, Funding acquisition, Methodology, Software, Visualization, Writing – original draft, Writing – review & editing), YS1 (Methodology, Investigation, Data curation, Writing – review & editing, Supervision), TI (Methodology, Investigation, Writing – review & editing, Supervision), KT (Methodology, Writing – review & editing), TO (Investigation, Resources, Writing – review & editing), RH (Investigation, Writing – review & editing), HO (Investigation, Resources, Writing – review & editing), YS2 (Conceptualization, Formal analysis, Methodology, Software, Writing – review & editing), MN (Conceptualization, Investigation, Resources, Funding acquisition, Writing – review & editing, Supervision, Project administration). All authors have read and approved the manuscript.

## Acknowledgements

We would like to thank Noriko Ishimura, Mika Kato, and Taku Sato for their work in participant recruitment and for their support in conducting the experiments, and Yukiko Yoshikane for her support with the experiments. We would like to thank Sayuri Takeda and Maiko Inoue for administering and scoring the clinical assessments. A portion of the TD sample was recruited through a website for recruiting research participants (https://www.jikken-baito.com). We also thank SOUKEN Co., Ltd. For its support with participant recruitment.

## Footnotes

1 These verbally described data were not analyzed here, as they were outside the scope of the present research question.

