## Supplementary Information for "Reduced EEG Complexity and Its Association with Social Communication in Adults with Autism Spectrum Disorder: A Multiscale Entropy Study"

**Naoe et al.**

### Supplementary Section 1

#### Additional data for main analyses 1

**Table S1 Experimental design.**

| Condition | Experiment | Stimuli | Time |
|---|---|---|---|
| Resting | Resting state with eyes closed. | - | 7 min |
| Movie Viewing 1 | Dynamically changing abstract visual patterns. | *Inscapes* | 7 min |
| Movie Viewing 2 | Animated character story with social-emotional scenes. | *Partly Cloudy* | 5 min |

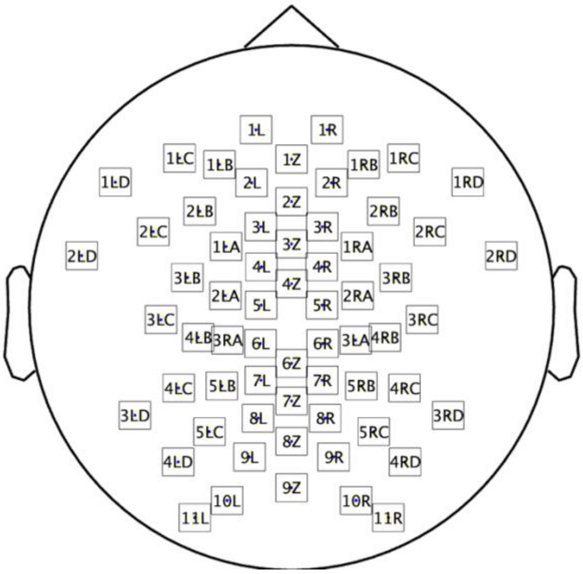


**Fig. S1 The equidistant scalp-position montage.**

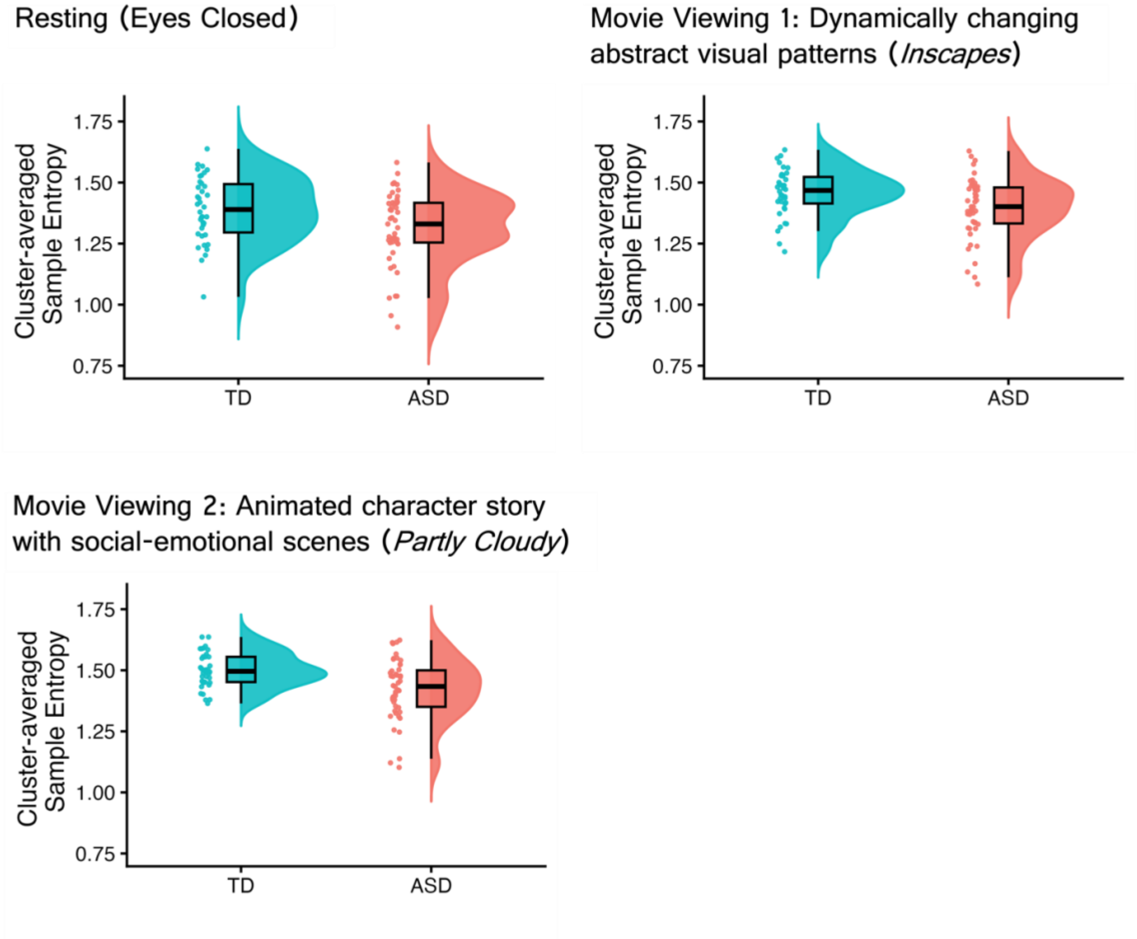


**Fig. S2 Distribution of the mean SampEn values.** Values are shown across conditions and are averaged over channels and scales within the cluster showing a significant group difference.

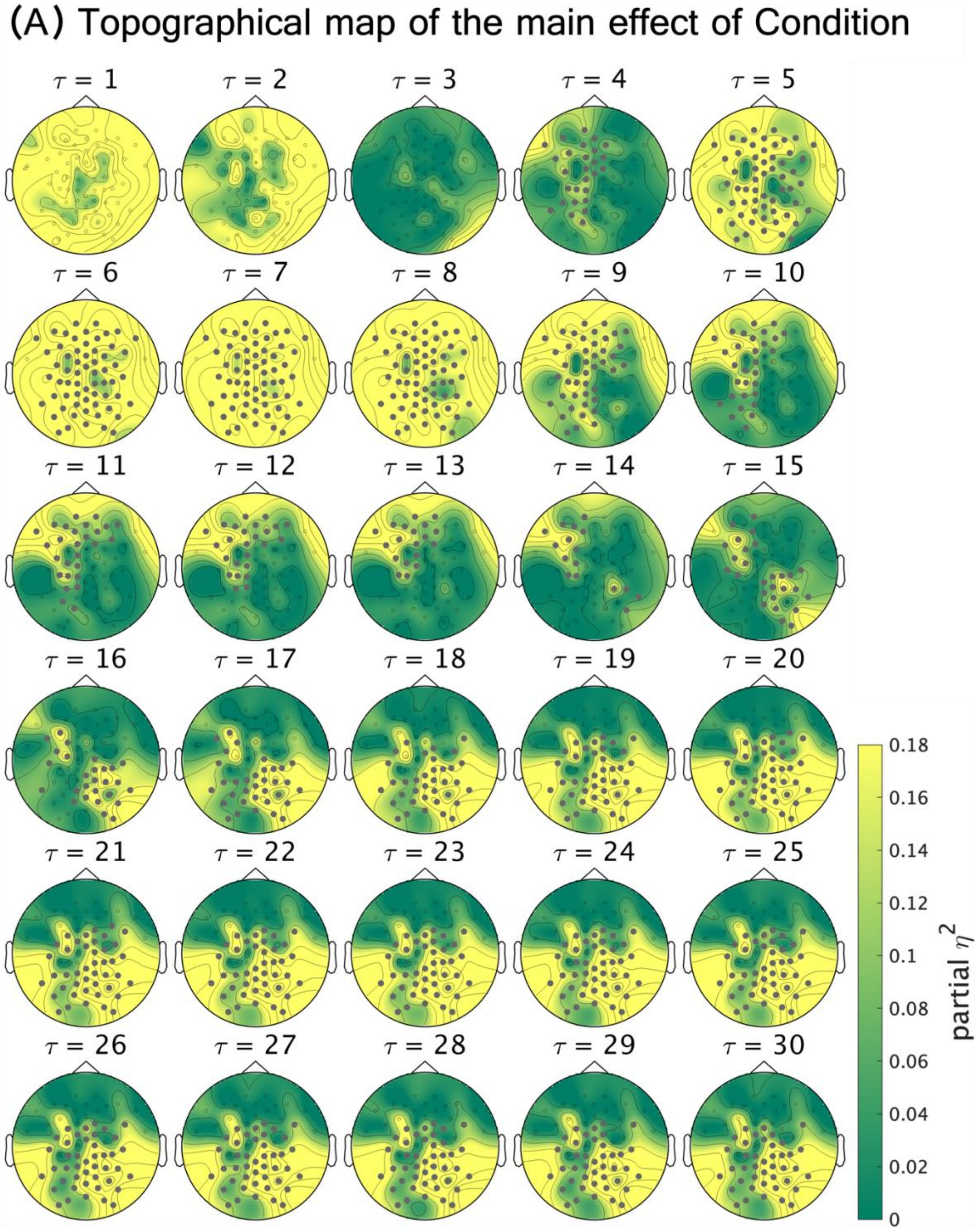


**Fig. S3 Differences in MSE across experimental conditions.** Topographical maps showing channel–scale partial $\eta^2$ effect-size maps for the Condition effect, derived from $F$ statistics computed from age- and sex-adjusted SampEn values. Each map represents condition-related SampEn effects at a given scale, corresponding to time series coarse-grained to 200/τ Hz, where τ denotes the scale factor. Channels belonging to the significant cluster are highlighted in gray. Statistical significance was assessed using one–sided cluster–based permutation tests (10,000 permutations; cluster-forming threshold $p < 0.05$; cluster–level $p < 0.05$).

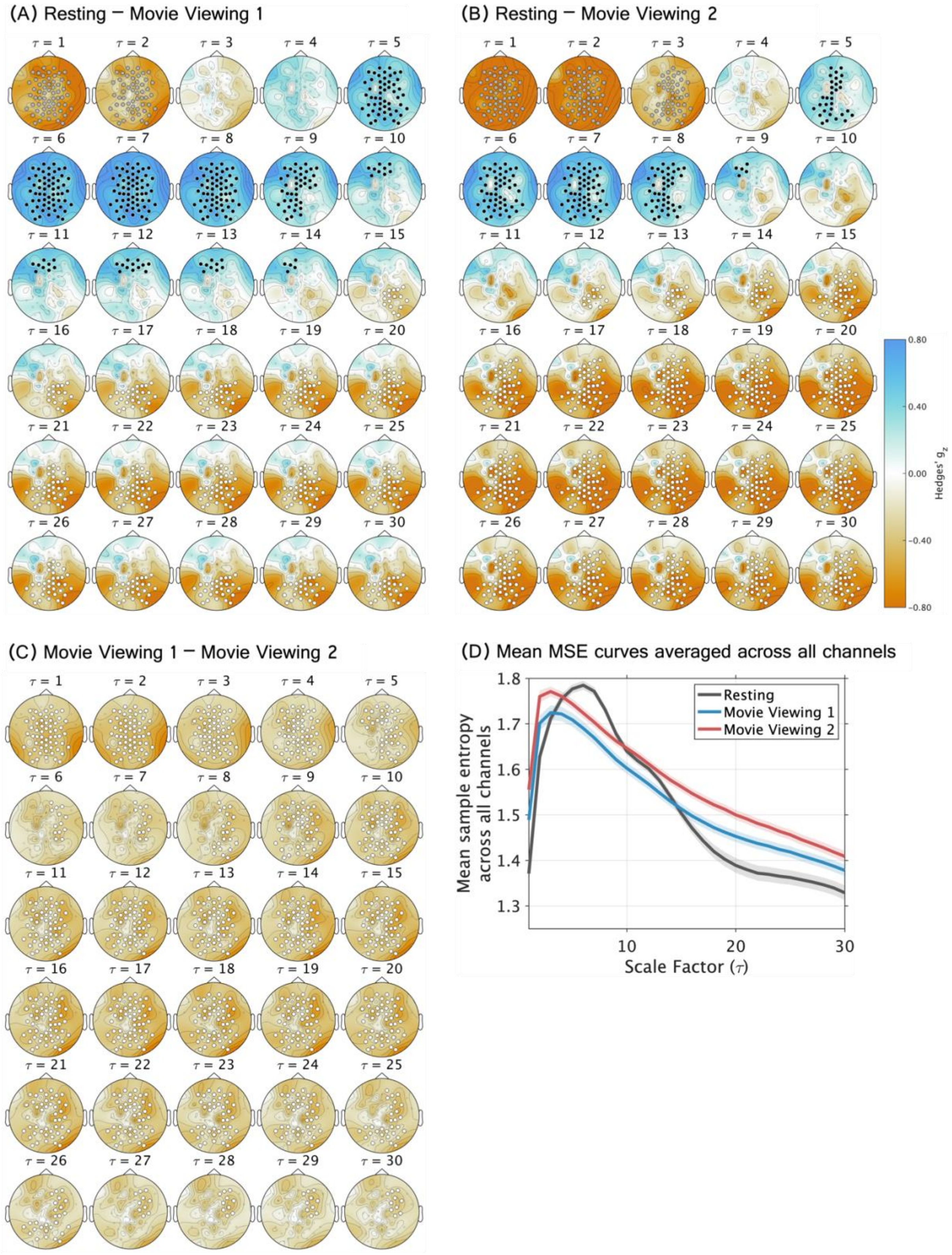


**Fig. S4 Pairwise comparisons of MSE across experimental conditions.** (A–C) Topographical channel–scale maps show Hedges' g for each between-condition contrast. At each channel and scale,

SampEn was computed from time series obtained by coarse-graining the original signal to an effective sampling rate of 200/τ Hz, where τ denotes the scale factor. A t statistic was calculated at each channel–scale point, and significance was assessed using two-sided cluster-based permutation tests (10,000 permutations; cluster-forming threshold, $p < 0.05$). Cluster-level p values were jointly FDR-corrected across the three pairwise comparisons. (A) Resting states with eyes closed vs Movie Viewing 1 (*Inscapes*). (B) Resting vs Movie Viewing 2 (*Partly Cloudy*). (C) Movie Viewing 1 (*Inscapes*) vs Movie Viewing 2 (*Partly Cloudy*). (D) Whole-scalp mean MSE curves and scale ranges of significant condition-difference clusters across experimental conditions. Line plots for each condition show mean MSE averaged across all channels (Resting: black, Movie Viewing 1: blue; Movie Viewing 2: red), with shaded bands indicating the 95% confidence intervals. The rectangles below indicate the scale ranges of clusters showing significant differences for each contrast (FDR-corrected cluster-level $ps < .05$). Different fill colors (gray, black, and white) denote distinct clusters within the same contrast and correspond to the cluster colors used in the topographical maps.

### Supplementary Section 2

**Sensitivity analyses for channel interpolation and montage harmonization**

The sensitivity-analysis sample comprised 33 ASD and 27 TD participants after the exclusion of 1 participant owing to a data-saving failure (Table S2). Among these participants, 24 ASD and 27 TD participants completed the WAIS-IV, and 5 ASD participants completed the WAIS-III. The remaining 4 ASD participants did not complete a WAIS assessment; however, their IQ estimates based on the JART were within the normal range (115.76, 105.65, 107.67, and 113.74).

**Table S2 Demographic information and clinical assessment scores of participants recorded with the equidistant scalp-position montage.**

| | ASD | TD | Group Difference |
|---|---|---|---|
| N | 33 (Female = 7) | 27 (Female = 5) | $\chi^2(1) = 0.00, p = 1.00$ |
| Age-years | 38.64±9.75 | 34.74±12.66 | $t = 1.31, p = .20$ |
| Estimated IQ (JART) | 109.88±8.59 | 111.79±6.88 | $t = –0.96, p = .34$ |
| FSIQ (WAIS-IV) | 108.54±16.29 (N = 24) | 117.41±14.61 (N = 27) | $t = –2.04, p < .05$ |
| VCI (WAIS-IV) | 111.75±16.49 (N = 24) | 117.67±15.74 (N = 27) | $t = –1.31, p = .20$ |
| PRI (WAIS-IV) | 107.00±14.46 (N = 24) | 113.11±15.55 (N = 27) | $t = –1.45, p = .15$ |

| | | | |
|---|---|---|---|
| WMI (WAIS-IV) | 107.42±14.79 (N = 24) | 115.56±16.39 (N = 27) | $t = –1.86$, $p = .07$ |
| PSI (WAIS-IV) | 96.08±15.55 (N = 24) | 108.07±13.76 (N = 27) | $t = –2.90$, $p < .01$ |
| FSIQ (WAIS-III) | 94.00±11.73 (N = 5) | - | - |
| VIQ (WAIS-III) | 98.00±11.90 (N = 5) | - | - |
| PIQ (WAIS-III) | 90.60±9.42 (N = 5) | - | - |
| ADOS-2 Total | 9.82±2.98 | 2.22±1.76 | $t = 12.27$, $p < .001$ |
| ADOS-2 SA | 8.30±2.43 | 2.07±1.62 | $t = 11.87$, $p < .001$ |
| ADOS-2 RRB | 1.52±1.03 | 0.15±0.36 | $t = 7.08$, $p < .001$ |

Note: Age and scores are presented as means and standard deviations. Sex distributions were compared using the Chi-square test. Continuous variables were compared using Welch's two-sample *t*-test. FSIQ, full-scale intelligence quotient; VCI, verbal comprehension index; PRI, perceptual reasoning index; WMI, working memory index; PSI, processing speed index; VIQ, verbal intelligence quotient; PIQ, performance intelligence quotient.

**Table S3 Number of participants with interpolation at each channel location. Only channels interpolated in at least one participant are shown.**

| Channel | ASD, n/N (%) | TD, n/N (%) |
|---|---|---|
| 1Z | 1/33 (3.0) | 0/27 (0.0) |
| 4Z | 1/33 (3.0) | 0/27 (0.0) |
| 6Z | 1/33 (3.0) | 0/27 (0.0) |
| 7Z | 2/33 (6.1) | 0/27 (0.0) |
| 1L | 1/33 (3.0) | 2/27 (7.4) |
| 6L | 2/33 (6.1) | 0/27 (0.0) |
| 9L | 1/33 (3.0) | 0/27 (0.0) |
| 11L | 3/33 (9.1) | 6/27 (22.2) |
| 1R | 1/33 (3.0) | 1/27 (3.7) |
| 4R | 1/33 (3.0) | 0/27 (0.0) |
| 5R | 4/33 (12.1) | 0/27 (0.0) |
| 6R | 1/33 (3.0) | 0/27 (0.0) |
| 7R | 1/33 (3.0) | 0/27 (0.0) |
| 9R | 1/33 (3.0) | 0/27 (0.0) |
| 11R | 4/33 (12.1) | 5/27 (18.5) |
| 1LB | 0/33 (0.0) | 1/27 (3.7) |
| 1LC | 0/33 (0.0) | 1/27 (3.7) |
| 3LD | 2/33 (6.1) | 0/27 (0.0) |

| Channel | ASD, n/N (%) | TD, n/N (%) |
| --- | --- | --- |
| 1RC | 1/33 (3.0) | 0/27 (0.0) |
| 1RD | 1/33 (3.0) | 0/27 (0.0) |
| 2RD | 0/33 (0.0) | 1/27 (3.7) |
| 3RD | 1/33 (3.0) | 1/27 (3.7) |
| 4RD | 0/33 (0.0) | 1/27 (3.7) |

**Group difference and effect of experimental condition (participants recorded with the equidistant scalp-position montage)**

A significant cluster for the main effect of Group was identified (cluster-level $p < .05$); this cluster indicated lower SampEn at middle-to-long scales ($\tau = 10$–$30$, coarse-grained time series with effective sampling rates of 20–6.67 Hz) in individuals with ASD than in TD controls (Fig. S5A). This cluster included channels showing effects with magnitudes ranging from small-to-large (partial $\eta^2$ = 0.067 to 0.186). Fig. S5B shows the mean MSE curves across the channels included in this cluster, and Fig. S5C shows the distribution of participant-level cluster-averaged MSE values.

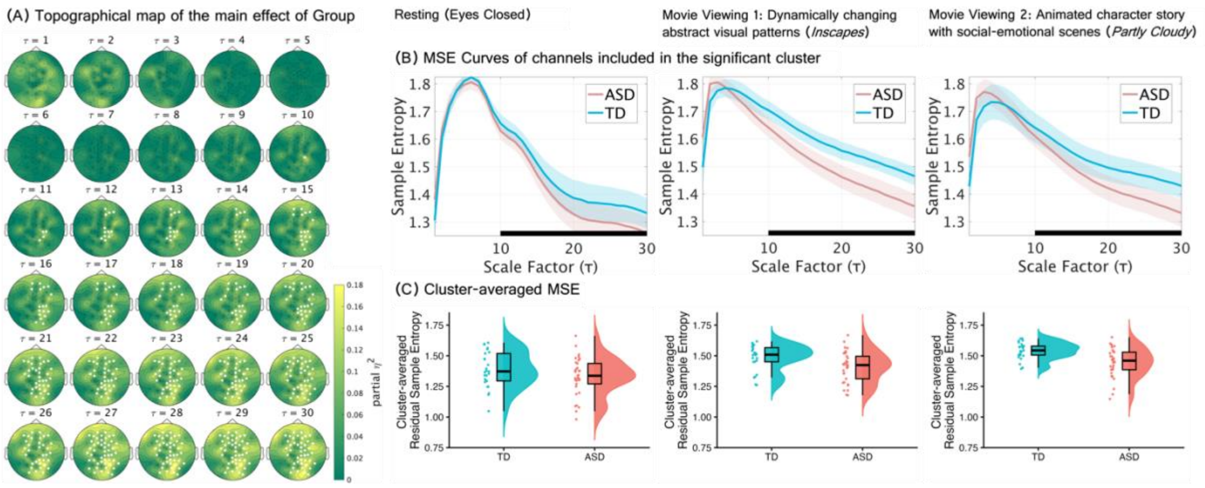


**Fig. S5 Group differences in EEG multiscale entropy (MSE) among participants recorded using an equidistant scalp-position montage.** (A) Topographical maps showing channel–scale partial $\eta^2$ effect-size maps for the Group effect, derived from $F$-statistics computed from age- and sex-adjusted SampEn values. Each topographical map represents the between-group differences in SampEn, computed from time series obtained by coarse–graining the original signal to 200/τ Hz. τ denotes the scale factor. A single significant cluster showing a Group effect was identified, and channels belonging to this cluster are highlighted in white. Statistical significance was assessed using one–sided cluster–based permutation tests (10,000 permutations; cluster-forming threshold $p < 0.05$; cluster–level $p < 0.05$). (B) Separate line plots for each condition show mean SampEn averaged across all channels included in the significant cluster at one or more scales (ASD: pink, TD: sky blue), with shaded bands indicating the 95% confidence intervals. In each plot, a black bar along the x-axis indicates the range of time scales spanned by the significant cluster. (C) Distribution of the mean SampEn values across conditions, averaged over channels and scales within the cluster showing significant group difference.

A significant main effect of Condition was identified in a cluster (cluster-level $p < .001$) spanning all scales ($\tau$ = 1–30, coarse-grained time series with effective sampling rates of 200–6.67 Hz), indicating that MSE varied across conditions irrespective of Group (Fig. S6). Partial $\eta^2$ values indicated small-to-large Condition effects across the cluster (partial $\eta^2$ = 0.052 to 0.663). Post hoc cluster-based permutation tests identified significant clusters for all three pairwise condition contrasts (FDR-corrected cluster-level *ps* < .05). Topographical maps showing the channels included in the significant clusters for each contrast, together with the corresponding effect sizes at each channel, are presented in Fig. S7A-C. As the significant clusters showed a widespread scalp distribution, mean MSE values averaged across all channels were plotted for each condition in Fig. S7D.

Resting versus Movie Viewing 1 and Resting versus Movie Viewing 2 showed broadly similar patterns, with three significant clusters identified in each contrast: both movie viewing conditions exhibited higher MSE than Resting at short scales (vs Movie Viewing 1: $\tau$ = 1–2; corresponding to coarse-grained time series with effective sampling rates of 200–100 Hz; FDR-corrected cluster-level p = .011; Hedges’ g = –1.057 to –0.258; vs Movie Viewing 2: $\tau$ = 1–3; corresponding to coarse-grained time series with effective sampling rates of 200–66.67 Hz; FDR-corrected cluster-level p = .010; Hedges’ g = –1.365 to –0.281) and long scales (vs Movie Viewing 1: $\tau$ = 15–30; corresponding to coarse-grained time series with effective sampling rates of 13.33–6.67 Hz; FDR-corrected cluster-level p = .003; Hedges’ g = –1.071 to –0.257; vs Movie Viewing 2: $\tau$ = 13–30; corresponding to coarse-grained time series with effective sampling rates of 15.38−6.67 Hz; FDR-corrected cluster-level $p = .001$; Hedges' $g = -1.499$ to $-0.257$), whereas MSE was higher during Resting at short-to-middle scales than both movie viewing conditions (vs Movie Viewing 1: $\tau = 4$–13; corresponding to coarse-grained time series with effective sampling rates of 50–15.38 Hz; FDR-corrected cluster-level $p = .005$; Hedges' $g = 0.260$ to 1.056; vs Movie Viewing 2: $\tau = 5$–9; corresponding to coarse-grained time series with effective sampling rates of 40–22.22 Hz; FDR-corrected cluster-level $p = .010$; Hedges' $g = 0.255$ to 0.862). The comparison between the two movie viewing conditions identified three distinct significant clusters spanning the full range of scales, all indicating higher MSE during Movie Viewing 2 than during Movie Viewing 1. The first cluster spanned $\tau = 1$–6 (corresponding to coarse-grained time series with effective sampling rates of 200–33.33 Hz; FDR-corrected cluster-level $p = .005$; Hedges' $g = -0.616$ to $-0.256$), the second spanned $\tau = 8$–25 (corresponding to coarse-grained time series with effective sampling rates of 25–8 Hz; FDR-corrected cluster-level $p = .003$; Hedges' $g = -0.840$ to $-0.255$), and the third spanned $\tau = 29$ (corresponding to coarse-grained time series with effective sampling rates of 6.90 Hz; FDR-corrected cluster-level $p = .044$; Hedges' $g = -0.431$ to $-0.262$).

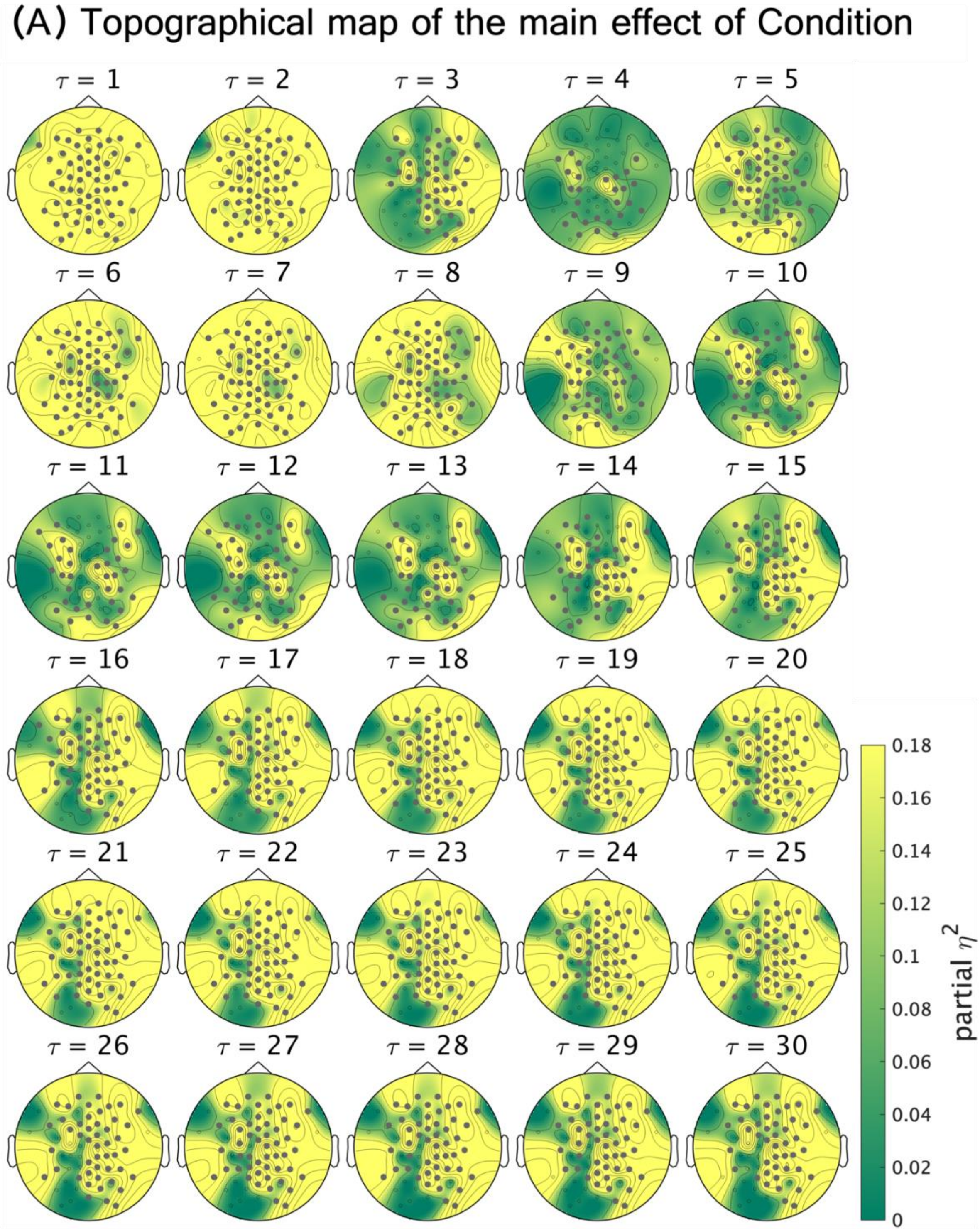


**Fig. S6 Differences in MSE across experimental conditions among participants recorded using an equidistant scalp-position montage.** Topographical maps showing channel–scale partial $\eta^2$ effect-size maps for the Condition effect, derived from $F$ statistics computed from age- and sex-adjusted SampEn values. Each map represents condition-related SampEn effects at a given scale, corresponding to time series coarse-grained to 200/τ Hz, where τ denotes the scale factor. Channels belonging to the significant cluster are highlighted in gray. Statistical significance was assessed using one–sided cluster–based permutation tests (10,000 permutations; cluster-forming threshold $p < 0.05$; cluster–level $p < 0.05$).

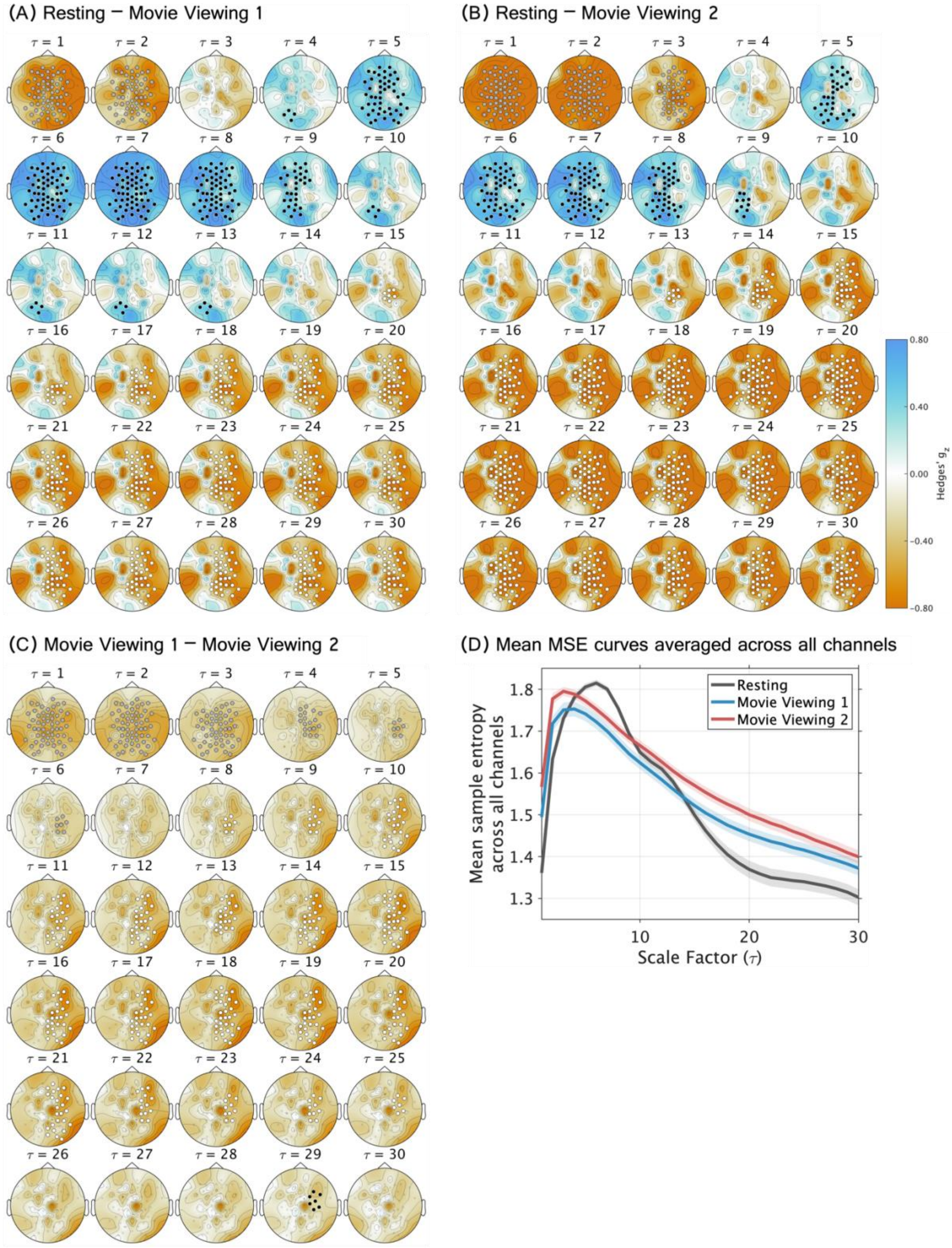


**Fig. S7 Pairwise comparisons of MSE across experimental conditions among participants recorded using an equidistant scalp-position montage.** (A–C) Topographical channel–scale maps show Hedges' g for each between-condition contrast. At each channel and scale, SampEn was computed from time series obtained by coarse-graining the original signal to an effective sampling rate of 200/τ Hz, where τ denotes the scale factor. A t statistic was calculated at each channel–scale point, and significance was assessed using two-sided cluster-based permutation tests (10,000 permutations; cluster-forming threshold, $p < 0.05$). Cluster-level p values were jointly FDR-corrected across the three pairwise comparisons. (A) Resting states with eyes closed vs Movie Viewing 1 (*Inscapes*). (B) Resting vs Movie Viewing 2 (*Partly Cloudy*). (C) Movie Viewing 1 (*Inscapes*) vs Movie Viewing 2 (*Partly Cloudy*). (D) Whole-scalp mean MSE curves and scale ranges of significant condition-difference clusters across experimental conditions. Line plots for each condition show mean MSE averaged across all channels (Resting: black, Movie Viewing 1: blue; Movie Viewing 2: red), with shaded bands indicating the 95% confidence intervals. The rectangles below indicate the scale ranges of clusters showing significant differences for each contrast (FDR-corrected cluster-level *ps* < .05). Different fill colors (gray, black, and white) denote distinct clusters within the same contrast and correspond to the cluster colors used in the topographical maps.

**Cluster-extent comparison across experimental conditions (Non-Social vs Social, participants recorded with the equidistant scalp-position montage)**

At the primary cluster-forming threshold ($\alpha = .05$), the largest suprathreshold cluster was 612 channel–scale points more extensive during Movie Viewing 2 than during Movie Viewing 1 (two-sided Monte Carlo $p = .004$, Fig. S8A). In the robustness analysis, the extent difference remained positive across cluster-forming thresholds from $\alpha = .10$ to .01, ranging from 180 to 612 channel–scale points (Fig. S8B). The observed differences also remained outside the corresponding covariance-matched Monte Carlo null distributions across the threshold sweep (two-sided Monte Carlo ps $\leq .021$).

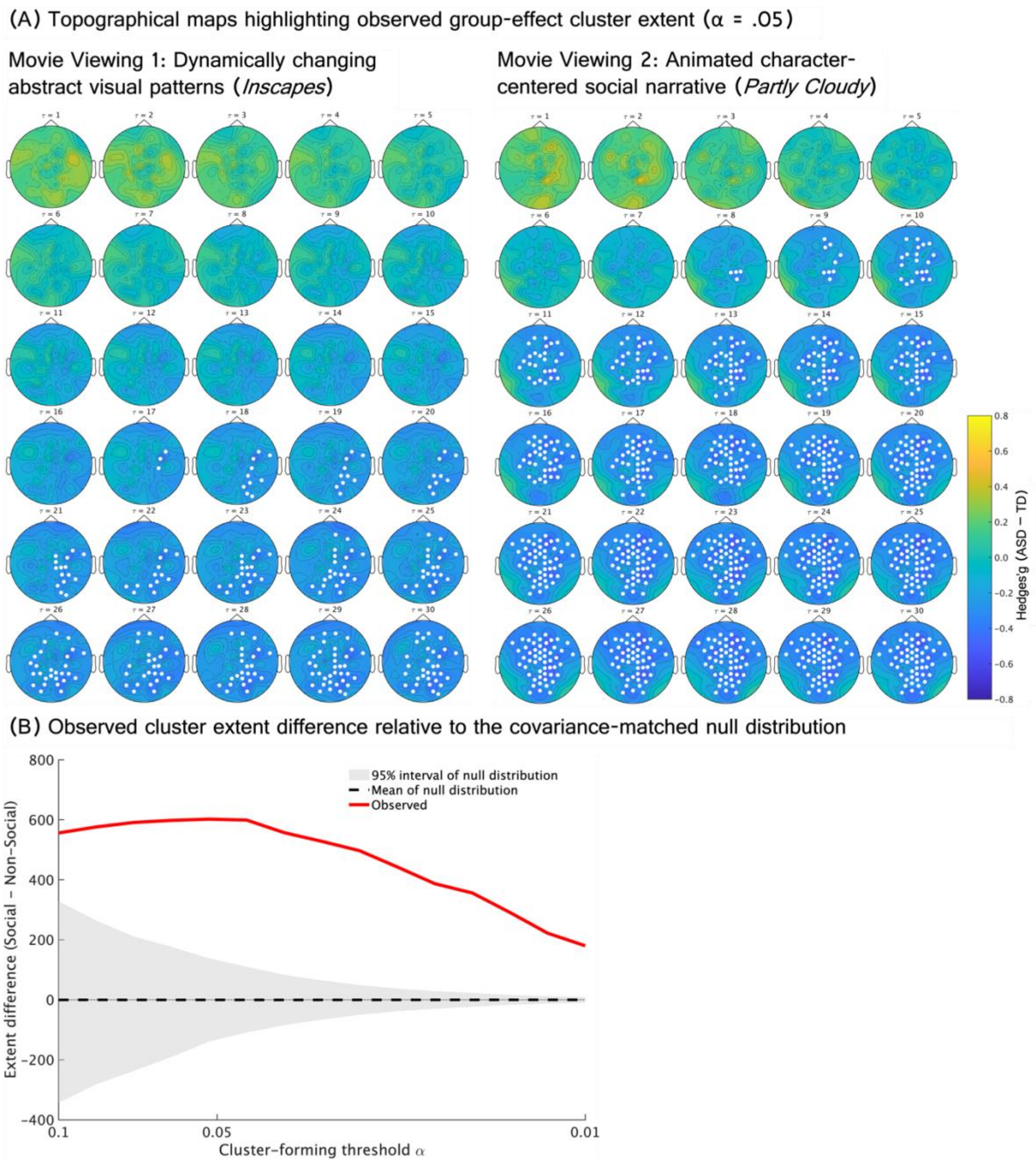


**Fig. S8 Differences in between-group cluster extent between movie viewing conditions in the equidistant-montage sample.** Movie Viewing 1 was compared with Movie Viewing 2. (A) Topographical channel–scale maps show Hedges' g (ASD–TD) for each Condition. Each topographical map represents the Hedges' g of between-group differences in SampEn, computed from time series obtained by coarse–graining the original signal to 200/$\tau$ Hz. $\tau$ denotes the scale factor. Channels belonging to the observed suprathreshold cluster are highlighted in white. (B) Observed cluster extent differences are shown across cluster-forming thresholds from $\alpha = 0.10$ to $\alpha = 0.01$. Cluster extent was quantified as the number of suprathreshold channel–scale points in the largest cluster. The covariance-matched null expectation was obtained from Monte Carlo Gaussian random fields preserving the empirical channel–scale covariance structure.

**Supplementary Section 3**

**Additional data for main analyses 2**

**Associations of LOSO-derived group-difference cluster MSE with ADOS-2 SA scores**

**Table S4 Results of the LME model examining the association between cluster-averaged MSE and ADOS-2 SA scores.**

| Fixed Effects | Estimate | 95% CI | | SE | t | p |
|---|---|---|---|---|---|---|
| (Intercept) | -0.000 | -0.026 | 0.026 | 0.013 | -0.010 | 0.992 |
| **ADOS-2 SA** | **-0.037** | **-0.071** | **-0.003** | **0.017** | **-2.177** | **0.032*** |
| Condition1 | 0.009 | -0.005 | 0.023 | 0.007 | 1.271 | 0.205 |
| **Condition2** | **0.046** | **0.032** | **0.060** | **0.007** | **6.406** | **< 0.001***** |
| Group | -0.045 | -0.090 | 0.001 | 0.023 | -1.954 | 0.054 |
| **age** | **-0.002** | **-0.004** | **-0.000** | **0.001** | **-2.025** | **0.046*** |
| sex | 0.010 | -0.040 | 0.059 | 0.025 | 0.392 | 0.696 |
| ADOS-2 SA : Condition1 | 0.018 | -0.001 | 0.037 | 0.010 | 1.916 | 0.057 |
| ADOS-2 SA : Condition2 | -0.011 | -0.029 | 0.008 | 0.010 | -1.113 | 0.267 |
| ADOS-2 SA : Group | 0.039 | -0.029 | 0.107 | 0.034 | 1.135 | 0.260 |

_Model: Cluster-averaged MSE ~ Group * ADOS-2 SA + Condition * ADOS-2 SA + age + sex + (1 | Participant)_

_Note_: ADOS-2: Autism Diagnostic Observation Schedule, Second Edition; SA: Social Affect. Condition1 (Resting: -1, Movie Viewing 1: 1, Movie Viewing 2: 0), Condition2 (Resting: -1, Movie Viewing 1: 0, Movie Viewing 2: 1). * p < .05, *** p < .001.

**Table S5 Results of the LME model examining the association between cluster-averaged MSE and ADOS-2 RRB scores.**

| Fixed Effects | Estimate | 95% CI | | SE | t | p |
|---|---|---|---|---|---|---|
| (Intercept) | 0.005 | -0.022 | 0.032 | 0.014 | 0.380 | 0.705 |
| ADOS-2 RRB | 0.013 | -0.033 | 0.060 | 0.023 | 0.577 | 0.566 |
| Condition1 | 0.010 | -0.004 | 0.024 | 0.007 | 1.371 | 0.172 |
| **Condition2** | **0.046** | **0.032** | **0.060** | **0.007** | **6.392** | **< 0.001*** ** |
| **Group** | **-0.066** | **-0.112** | **-0.020** | **0.023** | **-2.866** | **0.005** ** |
| age | -0.002 | -0.004 | 0.000 | 0.001 | -1.790 | 0.077 |
| sex | 0.006 | -0.047 | 0.058 | 0.026 | 0.210 | 0.834 |
| ADOS-2 RRB : Condition1 | -0.014 | -0.029 | 0.002 | 0.008 | -1.680 | 0.095 |

| Fixed Effects | Estimate | 95% CI | | SE | t | p |
|---|---|---|---|---|---|---|
| ADOS-2 RRB : Condition2 | -0.001 | -0.017 | 0.015 | 0.008 | -0.081 | 0.935 |
| ADOS-2 RRB : Group | -0.030 | -0.122 | 0.062 | 0.046 | -0.647 | 0.519 |

*Model: Cluster-averaged MSE ~ Group * ADOS-2 RRB + Condition * ADOS-2 RRB + age + sex + (1 | Participant)*

*Note*: ADOS-2: Autism Diagnostic Observation Schedule, Second Edition; RRB: Restricted and Repetitive Behaviors. Condition1 (Resting: -1, Movie Viewing 1: 1, Movie Viewing 2: 0), Condition2 (Resting: -1, Movie Viewing 1: 0, Movie Viewing 2: 1). ** p < .01, *** p < .001.

**Table S6 Results of the LME model examining the association between cluster-averaged MSE and ADOS-2 Total scores.**

| Fixed Effects | Estimate | 95% CI | | SE | t | p |
|---|---|---|---|---|---|---|
| (Intercept) | -0.002 | -0.029 | 0.025 | 0.014 | -0.120 | 0.905 |
| ADOS-2 Total | -0.035 | -0.072 | 0.001 | 0.018 | -1.924 | 0.058+ |
| Condition1 | 0.009 | -0.005 | 0.023 | 0.007 | 1.265 | 0.208 |
| **Condition2** | **0.046** | **0.032** | **0.060** | **0.007** | **6.369** | **< 0.001*** ** |
| Group | -0.045 | -0.091 | 0.002 | 0.023 | -1.912 | 0.059+ |

| Fixed Effects | Estimate | 95% CI | | SE | t | p |
|---|---|---|---|---|---|---|
| **age** | **-0.002** | **-0.005** | **-0.000** | **0.001** | **-2.031** | **0.046*** |
| sex | 0.007 | -0.044 | 0.058 | 0.026 | 0.286 | 0.775 |
| ADOS-2 Total : Condition1 | 0.010 | -0.008 | 0.029 | 0.009 | 1.107 | 0.270 |
| ADOS-2 Total : Condition2 | -0.009 | -0.028 | 0.010 | 0.009 | -0.946 | 0.345 |
| ADOS-2 Total : Group | 0.039 | -0.034 | 0.113 | 0.037 | 1.061 | 0.292 |

*Model: Cluster-averaged MSE ~ Group * ADOS-2 Total + Condition * ADOS-2 Total + age + sex + (1 | Participant)*

*Note*: ADOS-2: Autism Diagnostic Observation Schedule, Second Edition. Condition1 (Resting: -1, Movie Viewing 1: 1, Movie Viewing 2: 0), Condition2 (Resting: -1, Movie Viewing 1: 0, Movie Viewing 2: 1). + p < .10, * p < .05, *** p < .001.

**Event-related MSE changes during socially relevant and temporally matched non-social windows**

**Table S7 Empathic-pain and ToM events analyzed in the present study (adapted from Richardson et al. 2018)**

| Event | Event Number | Time | Duration (s) |
|---|---|---|---|
| ToM | T01 | 3:50–4:00 | 10 |
| | T02 | 2:36–2:52 | 16 |
| | T03 | 1:04–1:10 | 6 |
| | T04 | 4:32–4:46 | 14 |
| | T05 | 1:18–1:26 | 8 |
| | T06 | 3:38–3:48 | 10 |
| | T07 | 1:40–1:44 | 4 |
| Pain | P01 | 3:26–3:36 | 10 |
| | P02 | 1:56–2:14 | 18 |
| | P03 | 3:10–3:22 | 12 |
| | P04 | 0:50–0:56 | 6 |
| | P05 | 1:24–1:30 | 6 |

| | | |
|---|---|---|
| P06 | 4:00–4:10 | 10 |
| P07 | 4:52–4:56 | 4 |
| P10 | 1:08–1:12 | 4 |
| P12 | 2:52–2:56 | 4 |

Note: Event labels are those used by Richardson et al. (2018). We used windows identified in the first sample that showed at least partial temporal overlap with those independently identified in the second sample. Times are reported relative to the version used in the present study, from which the initial 10 s of the film were removed.

**Table S8 Results of the linear mixed-effects model on event-related MSE change.**

| Fixed Effects | Estimate | 95% CI | | SE | t | p |
|---|---|---|---|---|---|---|
| (Intercept) | 0.041 | -0.110 | 0.201 | 0.076 | 0.541 | 0.590 |
| Group (ASD/TD) | 0.051 | -0.170 | 0.289 | 0.111 | 0.459 | 0.647 |
| **Stimulus (Social/Non-Social)** | **-0.132** | **-0.190** | **-0.075** | **0.030** | **-4.428** | **< 0.001*** ** |
| Event (empathic pain/ToM) | -0.070 | -0.266 | 0.124 | 0.090 | -0.776 | 0.451 |
| ADOS-2 SA | -0.039 | -0.202 | 0.131 | 0.082 | -0.467 | 0.641 |
| **duration** | **-0.401** | **-0.501** | **-0.291** | **0.046** | **-8.652** | **< 0.001*** ** |

| Fixed Effects | Estimate | 95% CI | | SE | t | p |
|---|---|---|---|---|---|---|
| **sex** | **0.245** | **0.005** | **0.515** | **0.121** | **2.030** | **0.046*** |
| age | 0.024 | -0.082 | 0.133 | 0.054 | 0.455 | 0.650 |
| **Group : Stimulus** | **-0.166** | **-0.289** | **-0.046** | **0.063** | **-2.635** | **0.008**** |
| Group : Event | 0.011 | -0.113 | 0.133 | 0.063 | 0.169 | 0.866 |
| Stimulus : Event | 0.067 | -0.050 | 0.184 | 0.060 | 1.124 | 0.261 |
| Group : ADOS-2 SA | -0.122 | -0.453 | 0.227 | 0.166 | -0.735 | 0.465 |
| **Stimulus : ADOS-2 SA** | **-0.093** | **-0.174** | **-0.004** | **0.042** | **-2.231** | **0.026*** |
| Event : ADOS-2 SA | 0.027 | -0.055 | 0.108 | 0.042 | 0.640 | 0.522 |
| Group : Stimulus : Event | 0.053 | -0.193 | 0.312 | 0.126 | 0.421 | 0.674 |
| Stimulus : Event : ADOS-2 SA | -0.055 | -0.218 | 0.110 | 0.083 | -0.668 | 0.504 |

*Model: MSE change ~ Group * Stimulus * Event + Group * ADOS-2 SA + ADOS-2 SA * Stimulus * Event + duration + sex + age + (1 | Participant) + (1 | Time Window)*

*Note*: ADOS-2: Autism Diagnostic Observation Schedule, Second Edition; SA: Social Affect. * p < .05, ** p < .01, *** p < .001.

**The stability of MSE estimates**

To characterize the stability of MSE estimates obtained from EEG excerpts of different durations used in the event-related analysis, we performed a within-event resampling analysis. For each participant and each ToM or empathic-pain event window (4–18 s), the cluster-averaged MSE computed from the full event window served as the reference. For each target duration shorter than the full event (4, 6, 8, 10, 12, 14 or 16 s), we randomly selected up to 200 contiguous EEG excerpts with unique starting positions. These excerpts were therefore used to characterize within-event estimation variability rather than treated as independent observations. Because excerpts were sampled from the same event window, they could overlap, with the proportion of shared data generally increasing for longer target durations. MSE stability was therefore characterized from multiple complementary perspectives, including agreement with the full-event estimate, variability across excerpt positions, numerical estimability, and preservation of between-participant rank ordering. MSE was recalculated for each excerpt using the same parameters and participant-specific LOSO-derived mask as in the primary analysis. For each participant, event and target duration, agreement with the full-event reference was quantified using RMSE, and variability across excerpts was quantified as the SD of the cluster-averaged MSE estimates. Numerical estimability was assessed for each excerpt as the proportion of channel–scale points within the participant-specific mask that yielded finite MSE values; these proportions were then averaged across excerpts. Finally, for each event and target duration, Spearman's rank correlation across participants between the mean excerpt-based cluster MSE and the corresponding full-event reference

MSE was used to assess preservation of between-participant rank ordering.

RMSE and across-excerpt SD were highest for the shortest excerpts and generally decreased with increasing excerpt duration (Figs. S9 and S10). Spearman's rank correlations with the full-event reference estimates remained high across durations (Fig. S11, $\rho$ = 0.880–0.992), indicating that between-participant rank ordering was consistently preserved irrespective of duration. The proportion of channel–scale points yielding finite MSE values exceeded 98% for nearly all durations, although it was comparatively lower for the 4-s excerpts (Fig. S12).

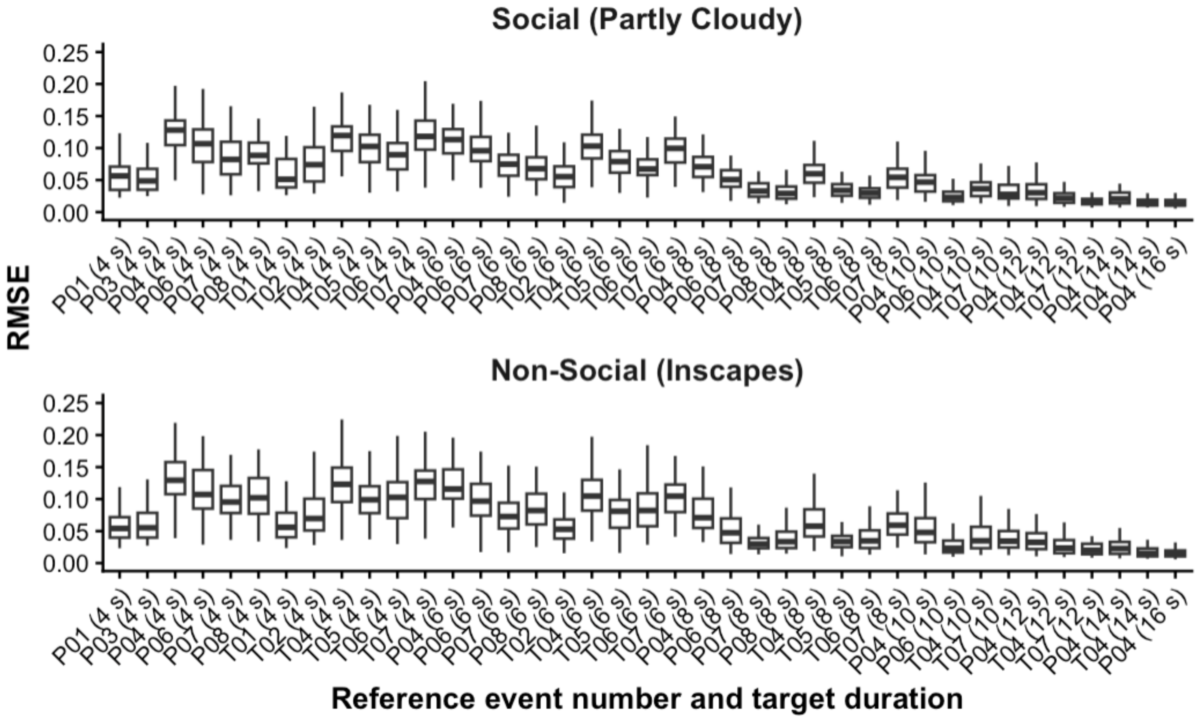


**Fig. S9 Root-mean-square error relative to the full-event MSE reference.** Each boxplot shows the distribution of participant-specific RMSE values for a given event and target duration. Event labels on the x-axis correspond to the event numbers listed in Table S7, with the Non-Social windows temporally matched to the corresponding Social events; values in parentheses indicate the target duration of the EEG excerpts extracted from each window.

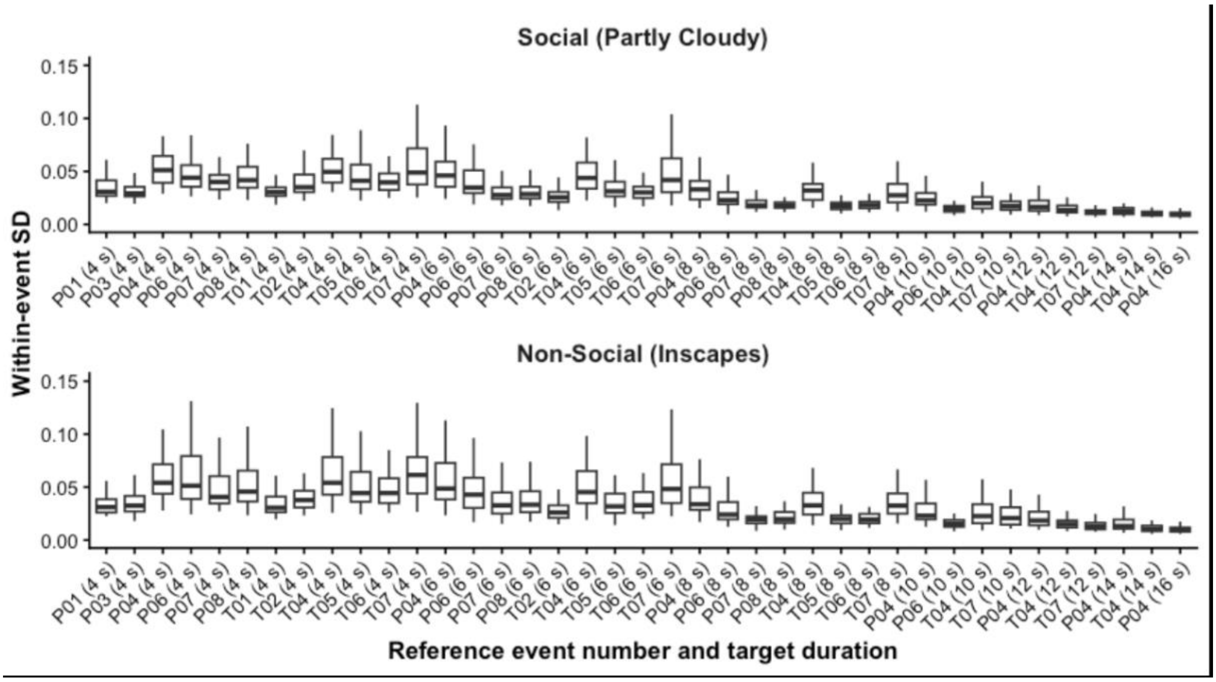


**Fig. S10 Within-event standard deviation of excerpt-based MSE estimates.** Each boxplot shows the distribution of participant-specific SD values for a given event and target duration. Event labels on the x-axis correspond to the event numbers listed in Table S7, with the Non-Social windows temporally matched to the corresponding Social events; values in parentheses indicate the target duration of the EEG excerpts extracted from each window. Lower SD indicates less variability in MSE estimates across excerpt positions within the event.

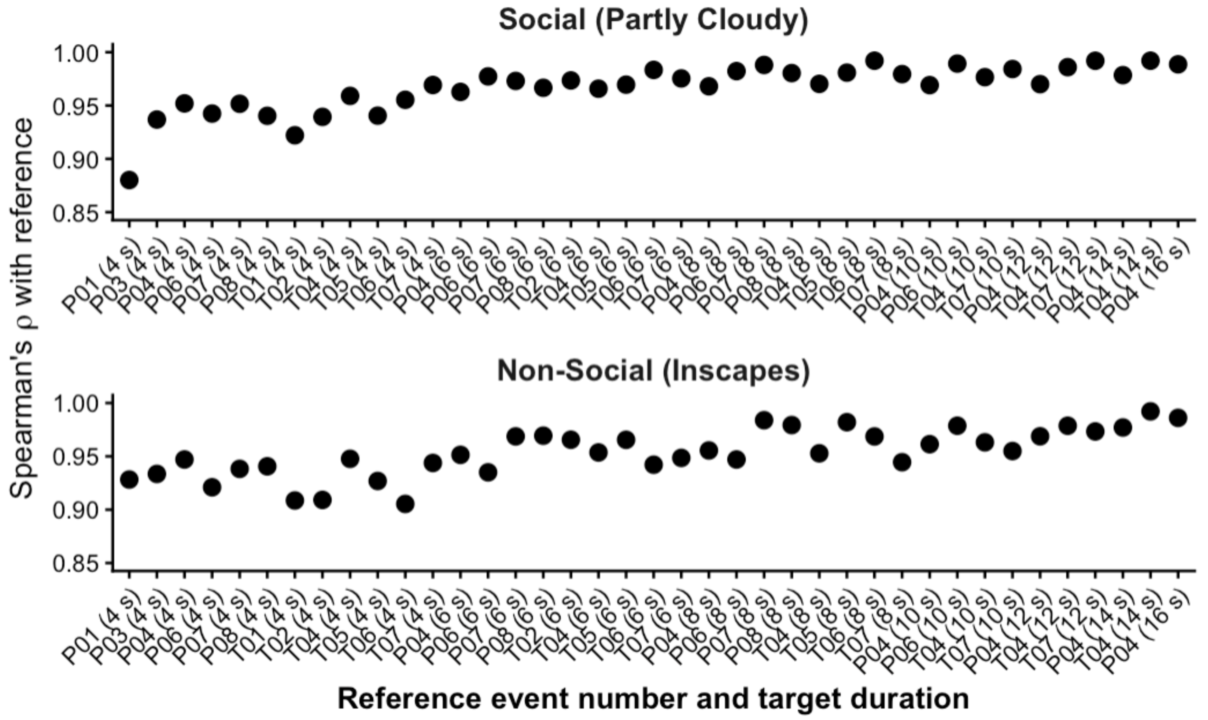


**Fig. S11 Preservation of between-participant rank ordering (Spearman's ρ).** For each event and target duration, Spearman's rank correlation was calculated across participants between the mean cluster-averaged MSE across the sampled EEG excerpts of that duration and the cluster-averaged MSE computed from the corresponding full event window. Event labels on the x-axis correspond to the event numbers listed in Table S7, with the Non-Social windows temporally matched to the corresponding Social events; values in parentheses indicate the target duration of the EEG excerpts extracted from each window.

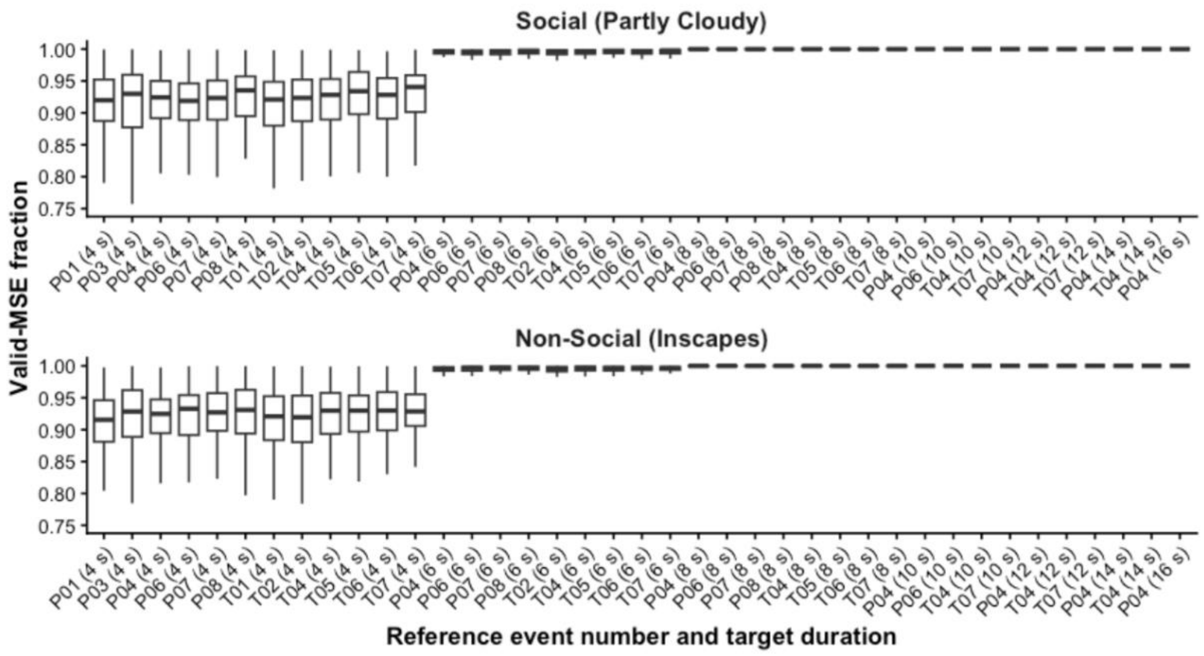


**Fig. S12 Proportion of channel–scale points yielding valid MSE estimates across excerpt durations.**

For each participant, event and target duration, the proportion of channel–scale points within the participant-specific leave-one-subject-out mask that yielded finite MSE values was calculated for each sampled EEG excerpt and then averaged across excerpts. Each boxplot shows the distribution of these participant-specific mean proportions for a given event and target duration. Event labels on the x-axis correspond to the event numbers listed in Table S7, with the Non-Social windows temporally matched to the corresponding Social events; values in parentheses indicate the target duration of the EEG excerpts extracted from each window.

**Table S9 Linear mixed-effects model of event-related MSE change with duration-dependent residual dispersion.**

| Fixed Effects | Estimate | 95% CI | | SE | z | p |
|---|---|---|---|---|---|---|
| (Intercept) | 0.041 | -0.110 | 0.191 | 0.077 | 0.530 | 0.596 |
| Group (ASD/TD) | 0.055 | -0.165 | 0.275 | 0.112 | 0.486 | 0.627 |
| **Stimulus (Social/Non-Social)** | **-0.142** | **-0.200** | **-0.085** | **0.029** | **-4.843** | **< 0.001*** ** |
| Event (empathic pain/ToM) | -0.072 | -0.250 | 0.107 | 0.091 | -0.786 | 0.432 |
| ADOS-2 SA | -0.037 | -0.200 | 0.126 | 0.083 | -0.445 | 0.656 |
| **duration** | **-0.399** | **-0.490** | **-0.308** | **0.047** | **-8.571** | **< 0.001*** ** |
| **sex** | **0.247** | **0.009** | **0.486** | **0.122** | **2.035** | **0.042*** |
| age | 0.024 | -0.082 | 0.130 | 0.054 | 0.446 | 0.656 |
| **Group : Stimulus** | **-0.154** | **-0.275** | **-0.033** | **0.062** | **-2.488** | **0.013*** |
| Group : Event | 0.011 | -0.110 | 0.133 | 0.062 | 0.184 | 0.854 |
| Stimulus : Event | 0.060 | -0.055 | 0.175 | 0.059 | 1.028 | 0.304 |
| Group : ADOS-2 SA | -0.131 | -0.459 | 0.198 | 0.168 | -0.778 | 0.436 |
| **Stimulus : ADOS-2 SA** | **-0.095** | **-0.175** | **-0.015** | **0.041** | **-2.324** | **0.020*** |
| Event : ADOS-2 SA | 0.023 | -0.057 | 0.103 | 0.041 | 0.562 | 0.574 |

| Fixed Effects | Estimate | 95% CI | | SE | z | p |
|---|---|---|---|---|---|---|
| Group : Stimulus : Event | 0.065 | -0.177 | 0.308 | 0.124 | 0.527 | 0.598 |
| Stimulus : Event : ADOS-2 SA | -0.043 | -0.203 | 0.117 | 0.082 | -0.529 | 0.597 |

*Model: MSE change ~ Group * Stimulus * Event + Group * ADOS-2 SA + ADOS-2 SA * Stimulus * Event + duration + sex + age + (1 | Participant) + (1 | Time Window); dispersion model: residual dispersion ~ duration.*

*Note*: ADOS-2: Autism Diagnostic Observation Schedule, Second Edition; SA: Social Affect. * p < .05, *** p < .001.

**Table S10 Linear mixed-effects model of event-related MSE change excluding 4-s windows.**

| Fixed Effects | Estimate | 95% CI | | SE | t | p |
|---|---|---|---|---|---|---|
| (Intercept) | 0.049 | -0.134 | 0.232 | 0.090 | 0.544 | 0.590 |
| Group (ASD/TD) | 0.071 | -0.155 | 0.297 | 0.114 | 0.625 | 0.534 |
| **Stimulus (Social/Non-Social)** | **-0.171** | **-0.240** | **-0.101** | **0.035** | **-4.824** | **< 0.001*** ** |
| Event (empathic pain/ToM) | -0.079 | -0.370 | 0.213 | 0.129 | -0.609 | 0.558 |
| ADOS-2 SA | -0.027 | -0.200 | 0.146 | 0.087 | -0.308 | 0.759 |
| **duration** | **-0.342** | **-0.494** | **-0.189** | **0.067** | **-5.076** | **0.001*** ** |

| Fixed Effects | Estimate | 95% CI | | SE | t | p |
|---|---|---|---|---|---|---|
| **sex** | **0.259** | **0.010** | **0.507** | **0.125** | **2.074** | **0.041*** |
| age | 0.019 | -0.091 | 0.129 | 0.055 | 0.347 | 0.730 |
| **Group : Stimulus** | **-0.149** | **-0.294** | **-0.004** | **0.074** | **-2.011** | **0.044*** |
| Group : Event | -0.044 | -0.189 | 0.102 | 0.074 | -0.588 | 0.557 |
| Stimulus : Event | 0.131 | -0.008 | 0.270 | 0.071 | 1.850 | 0.064+ |
| Group : ADOS-2 SA | -0.163 | -0.512 | 0.185 | 0.175 | -0.933 | 0.354 |
| **Stimulus : ADOS-2 SA** | **-0.109** | **-0.206** | **-0.012** | **0.049** | **-2.215** | **0.027*** |
| Event : ADOS-2 SA | 0.008 | -0.088 | 0.105 | 0.049 | 0.169 | 0.866 |
| Group : Stimulus : Event | 0.018 | -0.273 | 0.309 | 0.148 | 0.120 | 0.905 |
| Stimulus : Event : ADOS-2 SA | -0.027 | -0.220 | 0.166 | 0.098 | -0.270 | 0.787 |

*Model: MSE change ~ Group * Stimulus * Event + Group * ADOS-2 SA + ADOS-2 SA * Stimulus * Event + duration + sex + age + (1 | Participant) + (1 | Time Window)*

*Note*: ADOS-2: Autism Diagnostic Observation Schedule, Second Edition; SA: Social Affect. + $p < .10$, * $p < .05$, *** $p < .001$.
